# *De novo* Cyclic Peptides Allow Visualisation of the Monomeric and Functional Amyloid Conformations of the Kinase RIPK3

**DOI:** 10.1101/2025.11.05.686663

**Authors:** Jessica A. Buchanan, Olivia Lavidis, Nicholas Shields, Katriona Harrison, Huy T. Nguyen, Richard J. Payne, Toby Passioura, Megan Steain, Margaret Sunde

## Abstract

Receptor Interacting Protein Kinase 3 (RIPK3) is a central regulator of necroptosis and a key mediator of inflammatory signaling. Its function is orchestrated through RIP Homotypic Interaction Motif (RHIM)-dependent interactions with other RHIM-containing adapter proteins, forming amyloid-structured necrosomes that trigger kinase activation and subsequently lead to immunogenic cell lysis. Necroptosis eliminates infected or damaged cells and provokes a strong inflammatory response. Dysregulated necroptosis is implicated in chronic inflammatory diseases and associated with ischaemic injury. Despite separate structural elucidation of the isolated RIPK3 kinase domain and the amyloid fibrils formed by the RIPK3 RHIM-containing region, visualizing full-length human RIPK3 in live cells remains challenging due to a lack of specific tools. To address this, we employed Random Non-standard Peptide Integrated Discovery (RaPID) mRNA display to identify cyclic peptides that bind the RHIM-containing region of RIPK3 during its assembly into amyloid fibrils. Three peptides were selected for characterisation and demonstrate utility in visualizing RIPK3 in human cells. These peptides represent promising tools for probing RIPK3 localisation and modulating its function.

## Introduction

Receptor Interacting Protein Kinase-3 (RIPK3) is a crucial coordinating regulator of the lytic programmed cell death pathway known as necroptosis^[1-2]^. It also plays key roles in NF-κB signaling, inflammasome activation, and kinase-independent apoptosis^[1, 3-5]^. RIPK3 contains an 18–20-residue RIP Homotypic Interaction Motif (RHIM), through which it forms complexes called necrosomes with other RHIM-containing adapter proteins: RIPK1, TIR-domain-containing adapter-inducing interferon-β (TRIF), and Z-DNA binding-protein-1 (ZBP1)^[6]^. Necroptosis can be triggered by inflammatory mediators, the detection of pathogen-associated molecular patterns from fungal and bacterial infection, or through intracellular sensing of cellular or viral Z-form DNA or RNA^[7-8]^.

The role of RIPK3 as a hub within both protective inflammatory signalling and in cell death pathways, makes it of key interest in pathological conditions associated with unwanted necroptosis, as well as holding potential for therapeutic induction of necroptosis, for example, in cancer treatment^[5]^. RHIM sequences also occur in some viral proteins which inhibit host defence upon infection by incorporating into the necrosome complex via their RHIM and preventing necroptosis signal propagation^[9-13]^. Necrosomes have an amyloid cross-β fibril architecture, formed when the RHIMs from the constituent proteins interact to generate a β-sheet-rich fibril core that brings the adapter proteins into close contact with RIPK3^[14]^ (illustrated in Fig. 1).

**Figure 1.**
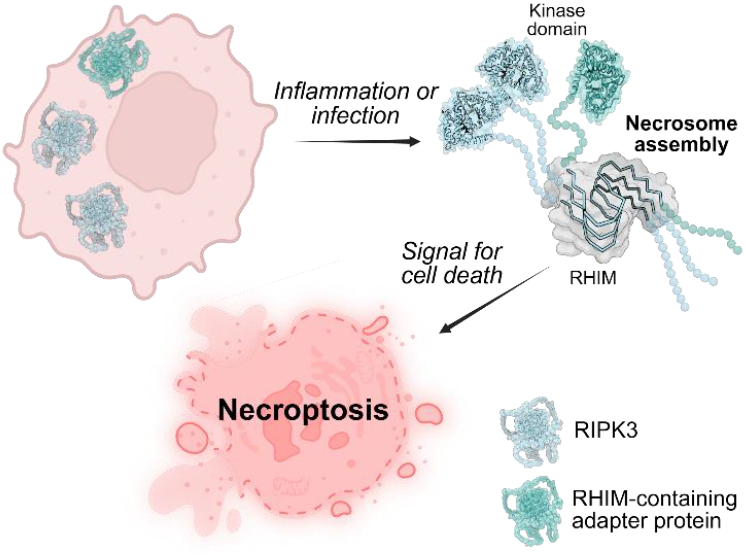
Schematic illustrating necrosome formation. Cytosolic, monomeric RIPK3 and RHIM-containing adapter proteins are activated by inflammation or fungal, bacterial or viral infection. RIPK3 and adapter proteins form the amyoid necrosome structure via their RHIMs, leaving kinase domains available along the fibril surface. Necrosome formation triggers downstream signalling for lytic, inflammatory cell death. Figure generated using BioRender.

The amyloid fibril-forming RHIM sequences render RIPK3 and the adapter proteins challenging candidates for structural studies. The structures of the isolated kinase domains of human and murine RIPK3 have been determined by crystallography^[15]^ and separately, the structures of the fibrils formed by the RHIM-containing regions of human and murine RIPK3 have been determined by cryoEM^[16]^. There is solution NMR evidence for a relatively structured “pre-RHIM” region of ∼20 residues immediately N-terminal of the RHIM^[17]^. However, little is known about the cellular location and interactions of full-length RIPK3 before and following induction of necroptosis. Efforts to express and study full-length RIPK3 have been unsuccessful, and even the isolated kinase domain is relatively unstable^[18]^. Studies of human RIPK3 (hRIPK3) have been hindered by a lack of effective tools to visualise the protein within live cells. Multiple antibodies have been raised against hRIPK3, but these suffer from poor selectivity, and they are ineffective as reporters in immunofluorescence-based studies for monitoring hRIPK3_[19-20]_.

Cyclic peptides offer opportunities for effective binding of proteins such as hRIPK3 which self-assemble into amyloid fibrils with large, iterative and hydrophobic surfaces. Indeed, the relatively large binding surface area of cyclic peptides makes them more selective than small molecule probes for such targets^[21]^. Additionally, because cyclic peptides require a structured target to bind to, target selective binding can be achieved by cyclic peptides in cases where selective antibodies cannot be identified^[22]^. The Random Non-standard Peptide Integrated Discovery (RaPID) system offers the combination of mRNA display with genetic code reprogramming, resulting in the ribosomal synthesis of expansive libraries containing >10^12^ macrocyclic peptides^[13]^.

Here we report the application of RaPID to identify and select novel cyclic peptide binders with affinity for the 387– 518 region of hRIPK3 (hRIPK3_387–518_), which is the RHIM-containing minimal region of human RIPK3 required for necrosome assembly^[23]^. The selection was applied during the process of amyloid fibril assembly. Three peptides were identified and their mode of binding to hRIPK3 characterised. Strikingly, these peptides were found to be capable of binding to hRIPK3 in both its monomeric and amyloid conformations. Additionally, these peptides are cell permeable and non-cytotoxic, allowing the visualisation of hRIPK3 within human cells and providing a starting point for the development of novel modulators of hRIPK3 activity.

## Results and Discussion

### Identification of Cyclic Peptide hRIPK3 Ligands using RaPID Screening

The minimum necrosome-forming region hRIPK3_387–518_ was first expressed with an N-terminal AviTag for biotinylation to allow for protein immobilisation to streptavidin beads during RaPID screening. This construct also contained a hexahistidine (His_6_) tag for purification and a ubiquitin fusion partner for improved expression (Fig. 2a). Complete biotinylation of this construct (henceforth referred to as BHUR3) was achieved via BirA ligase co-expression and biotin supplementation during protein expression. Fibril formation and fibrillar morphology was confirmed as equivalent to that of the non-biotinylated construct (HUR3) through monitoring of fibril formation kinetics using Thioflavin T (ThT), a widely used amyloid reporter fluorescent dye, and with transmission electron microscopy (Fig. S1).

**Figure 2.**
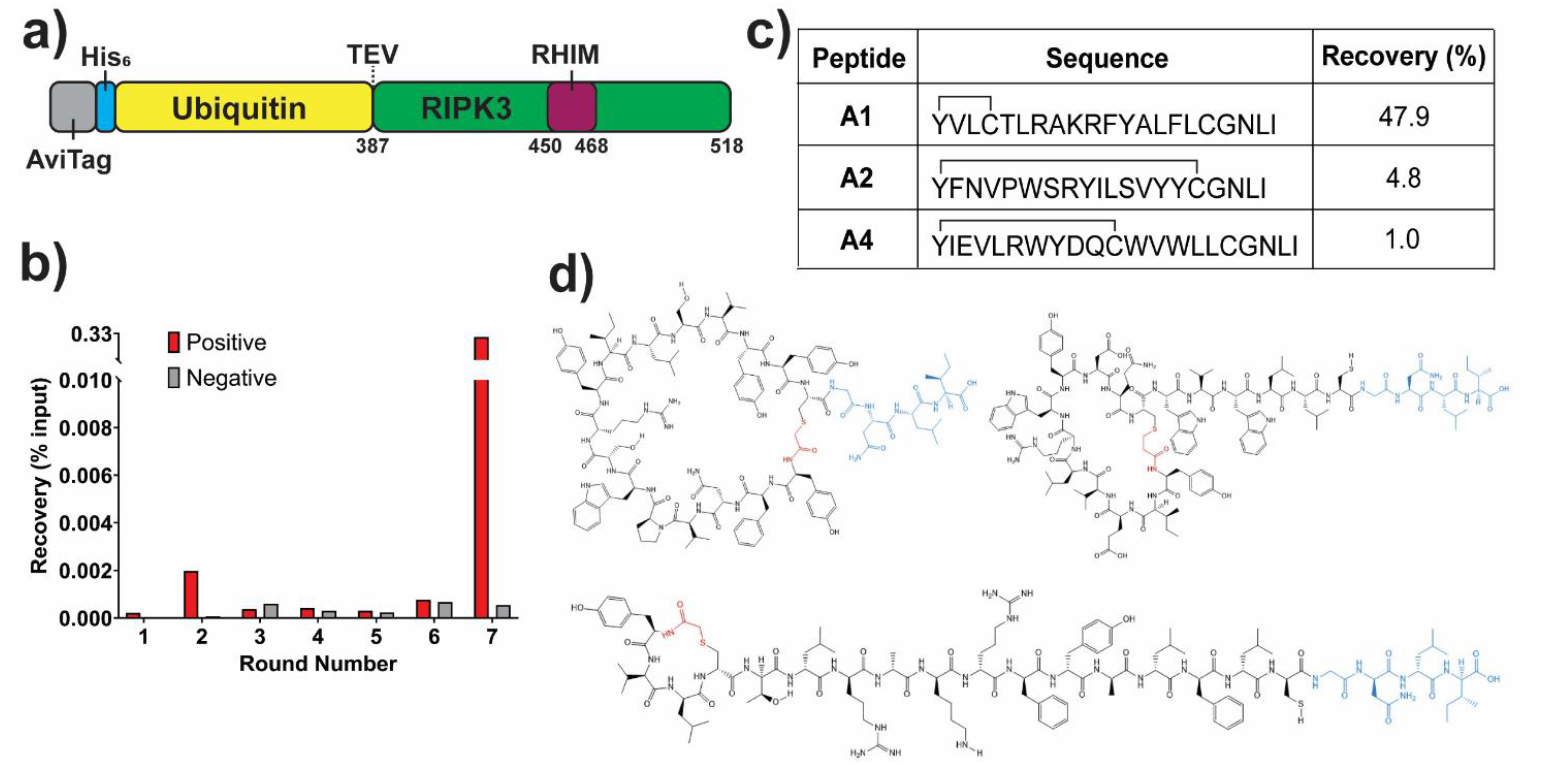
RaPID cyclic screening against BHUR3. a) Schematic of BHUR3 construct used for RaPID screening. b) Peptide recovery determined by RT-qPCR for positive and negative (counter) selection as a percentage of the total library input for each round of RaPID screening. Round 1 did not include a negative selection step. c) Sequences and total percent recovery of peptides selected for characterisation. d) Schematic of A1 (bottom), A2 (top left) and A4 (top right) cyclic peptides. GNLI linker shown in blue, cyclisation shown in red. Schematics created using ChemDraw.

For RaPID screening, a cyclic peptide library was produced via *in vitro* translation such that peptides consisted of an initiating *N*-chloroacetyl-L-tyrosine residue followed by a random sequence of 5–15 residues and concluding with a cysteine residue and C-terminal linker sequence (Gly-Asn-Leu-Ile). Spontaneous peptide cyclisation occurred during screening via thioether bond formation between the N-terminal chloroacetyl moiety and the closest cysteine sulfhydryl group. Peptides remained linked to their encoding nucleic acid through ligation of a puromycin molecule to the library of mRNA prior to translation^[13, 24]^.

RaPID screening was performed against BHUR3 during amyloid assembly. Briefly, BHUR3 was diluted out of denaturing buffer into the cyclic peptide library and allowed to incubate for 20 minutes at room temperature with gentle agitation, conditions known to lead to amyloid assembly. Following this incubation, the protein was immobilised onto streptavidin-coated magnetic beads to recover binding peptides. Counter-selection was performed against biotinylated His_6_-ubiquitin (BHU) to remove non-hRIPK3_387–518_-binding peptides from the library (see supporting information for full methodology). Enrichment was detected in the positive selection library after seven rounds of screening via RT-qPCR (Fig. 2b). Strikingly, almost 50% of the selected peptides corresponded to a single sequence, peptide A1, with the next most highly represented peptides A2 and A4 at 4.8 and 1.0%, respectively (Fig. 2c). These three peptides (A1, A2 and A4) were therefore synthesised by Fmoc-SPPS (see supporting information for synthetic details) and taken forward for characterisation.

### hRIPK3-Binding Cyclic Peptides Required Tetra-Lysine Tags to Abrogate Spontaneous Self-Assembly

The peptides were initially tested to determine whether they would influence amyloid assembly of hRIPK3, potentially acting as inhibitors if they preferentially bound to the monomeric form of hRIPK3 in a way that prevented RHIM:RHIM interactions. Unexpectedly, assay mixtures containing BHUR3 and any one of the peptides displayed a highly augmented ThT signal compared to BHUR3 alone (Fig. S2a–b), particularly peptide A1 (Fig. 3a), though there was negligible change in the rate of increase of ThT fluorescence. Further examination revealed that samples of peptide alone generated a strong ThT fluorescence signal (Fig. S2c–e). However, the peptide-only oligomers lacked the fibrillar morphology of amyloid fibrils formed by BHUR3, instead forming irregular clusters ∼50 nm in diameter (Fig. 3b–c). When the peptides were incubated with preformed hRIPK3_387–518_ fibrils, the size of the peptide oligomers was markedly reduced. Peptide species of 5−10 nm were observed decorating the length of the fibrils, indicative of preferential interaction with hRIPK3 fibrils over self-assembly (Fig. 3d, S3c–d).

**Figure 3.**
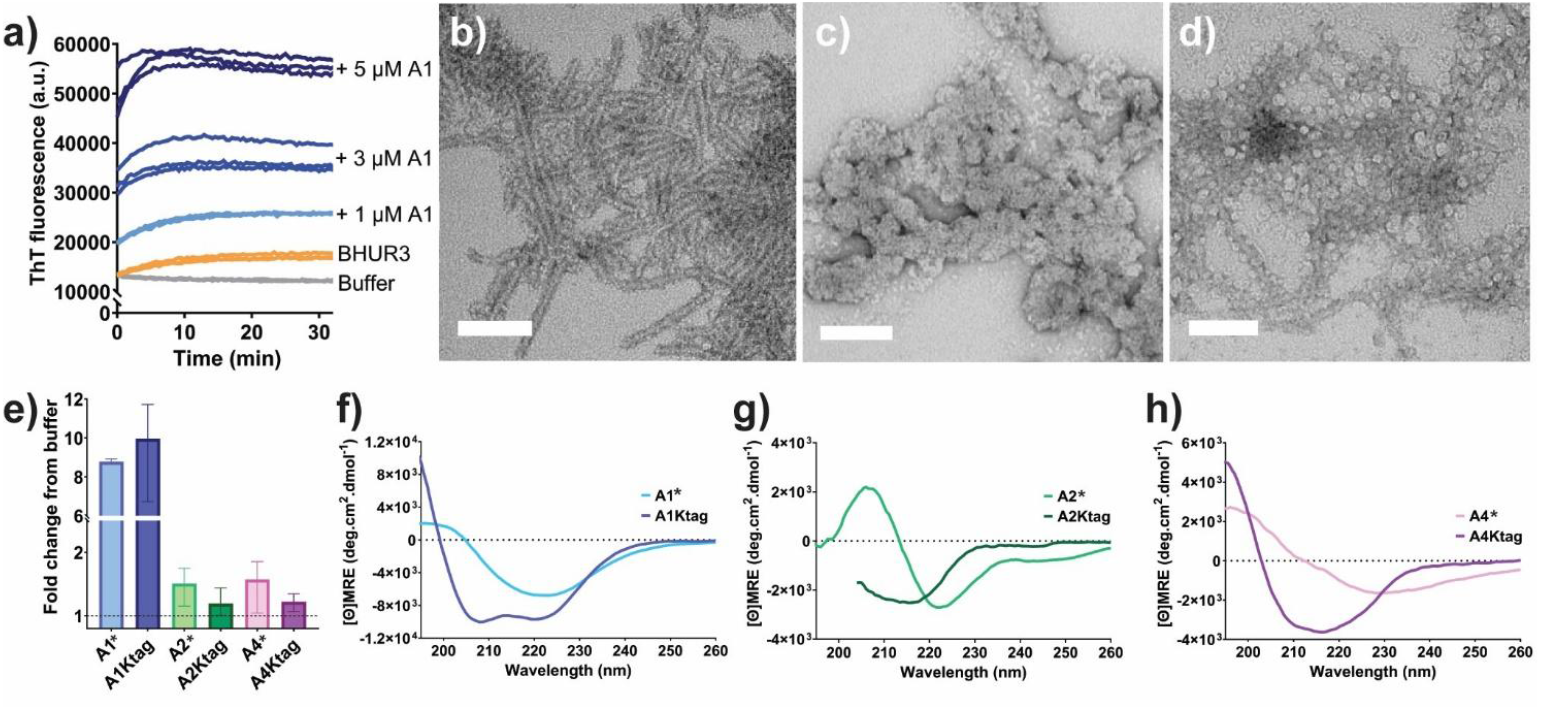
Influence of tetra-lysine tags on peptide aggregation propensity. a) ThT traces of 1 μM BHUR3 assembling with 0–5 μM A1 peptide in TBS buffer. b–d) Representative TEM images showing b) 0.2 mg/mL hRIPK3_387-518_ fibrils, c) 2 μM A1 peptide, and d) 0.2 mg/mL hRIPK3_387–518_ fibrils incubated with 2 μM A1 peptide. Images taken at 110 000X magnification, scale bars represent 100 nm. Grids were stained using 2% uranyl acetate. e) Average endpoint ThT signal after 1 hour assembly in TBS-T showing the fold change for peptide relative to buffer. Assay performed in triplicate, error bars represent range. f–h) CD spectra of original peptide constructs and their tetra-lysine-tagged counterparts using 0.3 mg/mL b) A1*/A1Ktag, c) A2*/A2Ktag, and d) A4*/A4Ktag. Peptides were solubilised in HFIP then diluted into 10 mM Tris pH 8.0, 0.01% Tween-20, data shows average of three collections. Data for A2Ktag truncated due to interfering buffer components <205 nm.

The addition of lysine residues to introduce charge-charge repulsion between peptide monomers is a strategy that has been successfully employed by others to improve cyclic peptide solubility without comprising binding^[21]^. Therefore, each peptide was resynthesised with either a single C-terminal lysine residue to facilitate conjugation of a sulfo-Cy5 moiety (A1*, A2* and A4*), or with a tetra-lysine tag in place of the C-terminal linker sequence (A1Ktag, A2Ktag, and A4Ktag). At the same time, the C-terminal cysteine in A1 and A4 was replaced by a serine to ensure correct cyclisation (see Table 1).

**Table 1.**
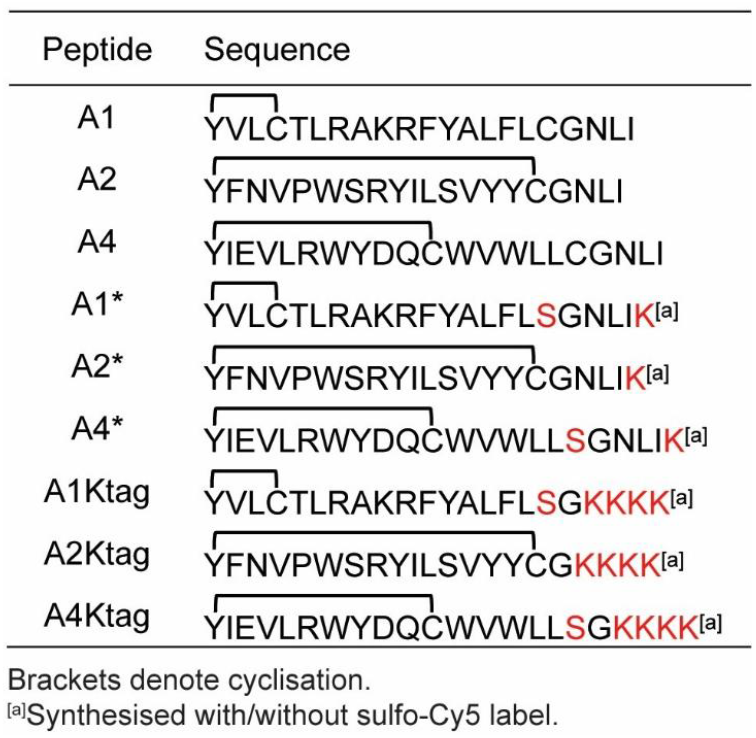

Single molecule photobleaching experiments demonstrated that with attachment of the sulfo-Cy5 label, the peptides were predominantly monomeric at 50 pM (Fig. S4). The introduction of the tetra-lysine sequence greatly reduced peptide self-assembly, as evidenced by reduction in ThT signal for A2Ktag and A4Ktag (Fig. 3e). Despite the increased solubility of A1Ktag compared to A1*, it still generated a strong ThT signal, possibly arising from an unusual interaction between this constrained cyclic peptide and the probe.

Circular dichroism (CD) spectropolarimetry was used to investigate the conformations of the peptides in solution. The untagged peptides did not display a strong β-signal at 215 nm, as would be expected for conventional amyloid structures that display ThT binding. Instead, they show minima between 220 and 230 nm, with some aggregation observed during the collection of spectra (Fig. 3f–h). All of the tetra-lysine-tagged peptides underwent a large conformation change and an increase in α-helicity resulting from the addition of the tetra-lysine tag. The observed increase in signal intensity was also indicative of reduced aggregation propensity. The tetra-lysine tagged variants, with or without the addition of a sulfo-Cy5 label, were therefore taken forward for all subsequent experiments.

### Identified Cyclic Peptides Bind hRIPK3 with Micromolar Affinity

TIRF fluorescence microscopy demonstrated binding of the sulfo-Cy5-labelled tetra-lysine-tagged peptides to hRIPK3_387-518_ fibrils immobilised on slides (Fig. 4a). Subsequently, these peptides were used to determine affinities for hRIPK3_387-518_ fibrils using microscale thermophoresis (MST). All displayed low micromolar affinity, calculated on the starting hRIPK3_387–518_ monomer concentration, with dissociation constant (*K*_*d*_) values of 1.96 ± 0.27, 1.26 ± 0.33 and 2.30 ± 0.39 μM for A1Ktag, A2Ktag and A4Ktag respectively (Fig. 4b). This analysis presumes a 1:1 binding mode; however, this stoichiometry is unlikely due to the self-assembling nature of both the peptides and their amyloid target. Given the relatively large size of the cyclic peptides, the binding site may span more than one hRIPK3 layer of the fibril.

**Figure 4.**
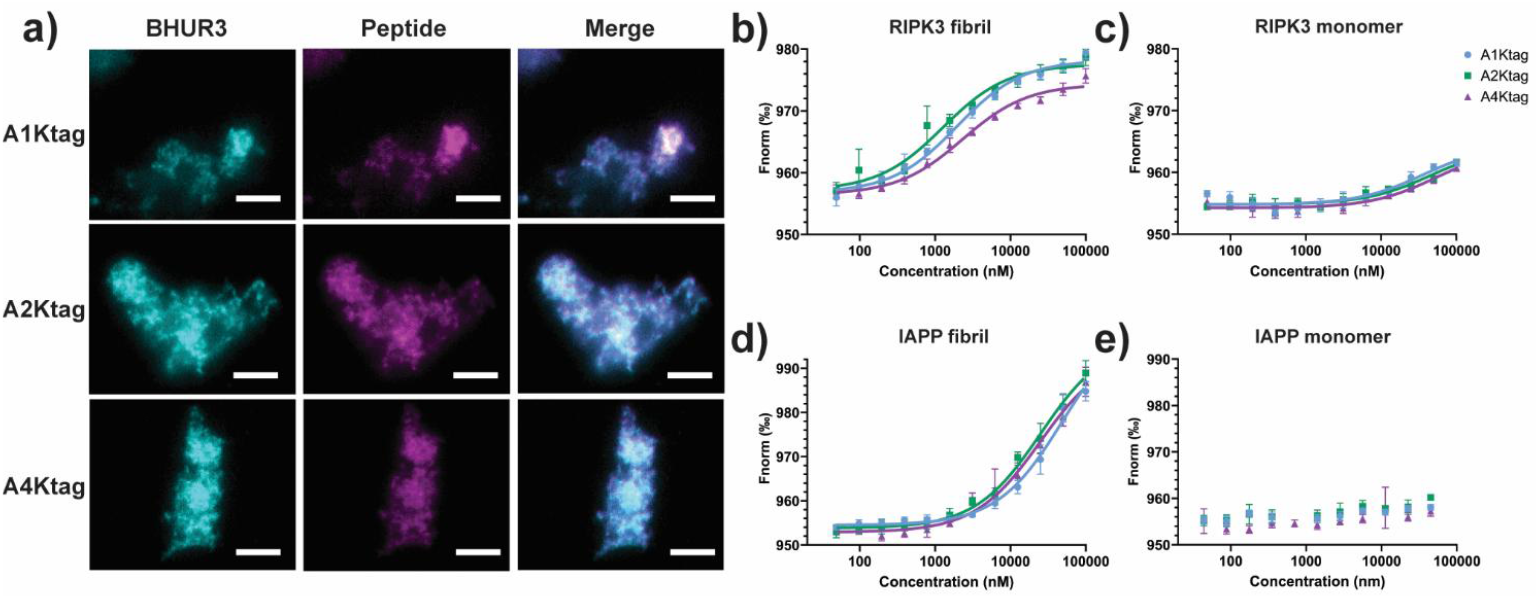
Binding of tetra-lysine-tagged peptides to hRIPK3_387–518_. a) TIRF images of 500 pM sulfo-Cy5-lablled peptides with preformed BHUR3 fibrils. Fibrils were adhered to a poly-L lysine-treated coverslip, then peptides diluted into TBS-T and incubated with fibrils for 15 minutes. Coverslips were washed thrice with TBS-T before imaging. Images taken at 100X magnification, scale bars represents 5 μm. b–e) MST traces of 20 nM tetra-lysine-tagged peptides with b) hRIPK3_387–518_ fibrils, c) HUR3 monomer, d) IAPP fibrils, or e) IAPP monomer. b), d) and e) were performed in TBS-T pH 8.0, 0.1 mg/mL BSA, 0.04% glycerol, 5% DMSO. c) was performed in 4 M urea, 20 mM Tris pH 8.0, 5% DMSO to maintain HUR3 in a monomeric conformation.

To investigate selectivity, the peptides were exposed to preformed human islet amyloid polypeptide (IAPP) or insulin fibrils, and to monomeric forms of hRIPK3_387-518_, IAPP, insulin and hen egg white lysozyme. The hRIPK3_387-518_ construct is extremely amyloidogenic and could only be maintained in a monomeric form at pH 8.0 in the presence of 4 M urea (Fig. S5a). Strikingly, some binding to monomeric hRIPK3_387–518_ was observed, albeit weak (>35 μM), while no binding to monomeric IAPP, insulin or lysozyme was detected (Fig. 4c–d and Fig. S5b, d). The affinity of the peptides for monomeric hRIPK3 under physiological conditions is likely to be higher than measured here, since hydrophobic interactions and H-bonding interactions are disfavoured in the presence of the denaturant. Some interaction with IAPP fibrils (Fig. 4d) and insulin fibrils (Fig. S5c) was detected, though these did not achieve saturation at the concentrations used for hRIPK3_387–518_ fibrils, indicating an affinity for these fibrils of likely >10 μM (all calculated *K*_*d*_ values available in Supplementary Table 1).

The results suggest that these peptides bind with high affinity to hRIPK3 and additionally interact with the cross-β structure common to all amyloid fibrils. This RaPID selection was performed during amyloid formation, and these results are consistent with the species of hRIPK3 populated during the selection, namely monomer, small oligomers and mature fibrils with a cross-β structure. In another study, peptides were identified from a RaPID screen against mature α-synuclein fibrils^[25]^. Those peptides exhibited dissociation constants of ∼3–10 μM and displayed the ability to induce liquid-liquid phase separation in α-synuclein, and under some conditions accelerated amyloid fibril formation^[25]^. Similar to the hRIPK3-binding peptides, not all identified peptides were specific for α-synuclein, highlighting that self-assembling proteins share similar structural features or properties, from the “stickers” of LLPS-prone proteins to the grooves present on cross-β amyloid fibrils.

### hRIPK3-Binding Peptides are Non-Toxic, Cell-Permeable and Reveal the Location of hRIPK3 in Cells undergoing Necroptosis

HT-29 cells provide a well-established system to model necroptotic cell death, as they express the cellular machinery required for apoptosis and necroptosis^[23, 26-27]^. This cell line was therefore selected to determine if the peptides could be used to visualise the localisation of hRIPK3 following induction of these cell death pathways. First, peptides were delivered after permeabilisation of the cell membrane. HT-29 cells were treated with vehicle

(DMSO, untreated control) or with TNF and the SMAC mimetic BV-6 to trigger apoptosis, or TNF, BV-6 and the pan-caspase inhibitor z-VAD to trigger necroptosis. Following a 6-hour incubation, cells were fixed, permeabilised and fluorescently labelled peptides were applied at 10 nM and then cells were extensively washed to remove unbound or weakly bound material. All three hRIPK3-binding peptides were able to enter the permeabilised cells (Fig. 5a). The A1Ktag, A2Ktag and A4Ktag peptides were each observed inside all cells, regardless of the treatment condition. This contrasts with the lack of uptake or binding of a peptide identified by a RaPID screen against an unrelated protein not expected to be present in HT-29 cells, which was also tested and displayed no binding in these cells under any condition. Cells exposed to the control peptide labelled with an Alexa Fluor™ 594 fluorophore were indistinguishable from cells not exposed to any sulfo-Cy5-labelled peptides. Together, these results indicate that the observed localisation within cells is related to specific hRIPK3 binding.

**Figure 5.**
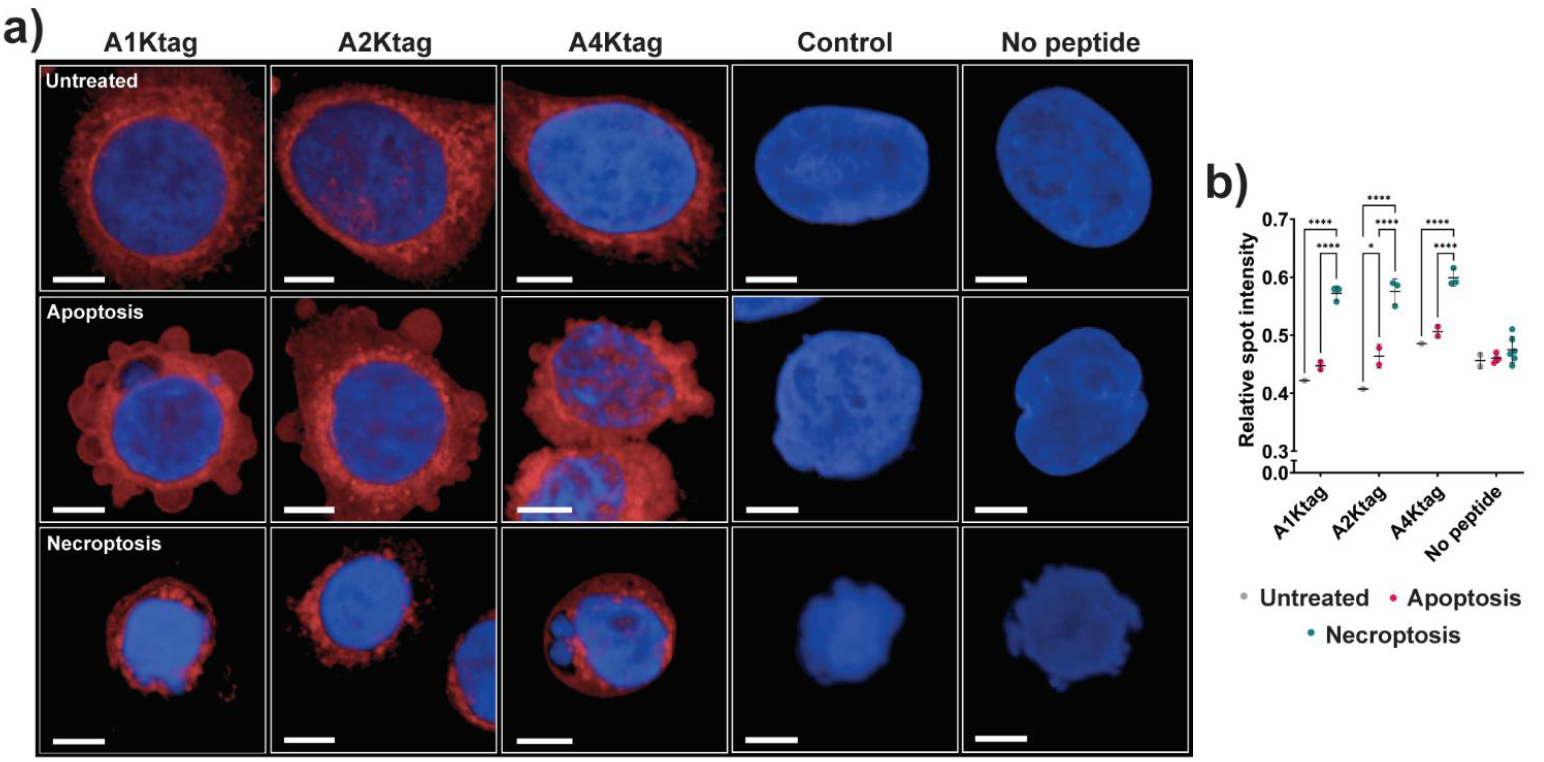
IFA using sulfo-Cy5 labelled tetra-lysine-tagged peptides in HT-29 cells. a) Confocal images of HT-29 cells treated with DMSO vehicle (Untreated), TNFα and BV-6 (Apoptosis) or TNFα, BV-6 and zVAD-fmk (Necroptosis), fixed, permeabilised then incubated with 10 nM sulfo-Cy5-labelled peptides (red) and DAPI (blue). Control peptide labelled with Alexa Fluor™ 594 and imaged using appropriate excitation wavelenth. Images collected at 63X magnification, scale bar represents 10 μm. b) Mean relative spot intensity within cells of each well as a ratio of spot intensity to background intensity. Mean and SD shown, a single biological replicate was performed, DMSO vehicle had single well, apoptosis duplicate wells and necroptosis triplicate wells for each peptide treatment. 2-way ANOVA using Tukey’s multiple comparisons test conducted for statistical analysis (* <0.05, ** <0.01, *** <0.001, **** <0.0001). Non-significant comparisons not indicated.

The hRIPK3-binding peptides in untreated and apoptosis-induced cells were predominantly dispersed throughout the cytoplasm. Some perinuclear concentration was observed, consistent with studies of murine RIPK3 and imaging of hRIPK3-associated proteins ^[20, 27]^. The distribution of the peptides highlighted the characteristic blebbing morphology associated with cells undergoing apoptosis. In contrast, in the cells undergoing necroptosis, the peptides were concentrated into puncta which occupied most of the cytoplasm, consistent with the formation of hRIPK3-rich necrosomes. Analysis of the intensity of the Cy5 signal in observed puncta within cells demonstrated that the relative spot intensity in cells undergoing necroptosis was significantly higher than in untreated cells in cells undergoing apoptosis (Fig. 5b). Notably, strong binding was observed even with the use of nanomolar concentrations of the peptides, suggesting that the true affinity for full-length hRIPK3 may be significantly higher than the low micromolar affinity measured *in vitro* against fibrils formed from hRIPK3_387-518_.

We then sought to assess the cell permeability and uptake of tetra-lysine-tagged peptides when incubated with live HT-29 cells. Peptide cell association with live cells was assessed via flow cytometry after a 4-hour incubation of peptides with cells in media. Flow cytometry showed high levels of peptide association compared to free Cy5 (Fig. 6a). We determined that the three hRIPK3-binding peptides are also not cytotoxic to live cells. HT-29 cells were incubated with the sulfo-Cy5-labelled peptides in media for 10 hours and no cell death was detected, as with no peptide (vehicle) controls (Fig. 6b). These data contrasted with cells treated with TSZ to induce necroptosis, which showed ∼40% cell death 8 hours post-treatment. The lack of cytotoxicity encouraged us to investigate peptide uptake into live cells further. Confocal microscopy was used to determine whether the peptides were taken up into the cells and their subcellular distributions. All three hRIPK3-binding peptides showed cellular uptake, with the majority of each peptide located within the cytoplasm (Fig. 6c). Some peptide was observed concentrated in organelles (e.g. endosomes), and a small amount was identified within the nucleus (Fig. 6c–d). Future work is required to confirm the uptake pathways, but these results indicate that the hRIPK3-binding peptides could be used to track necrosome assembly and fate in live cells.

**Figure 6.**
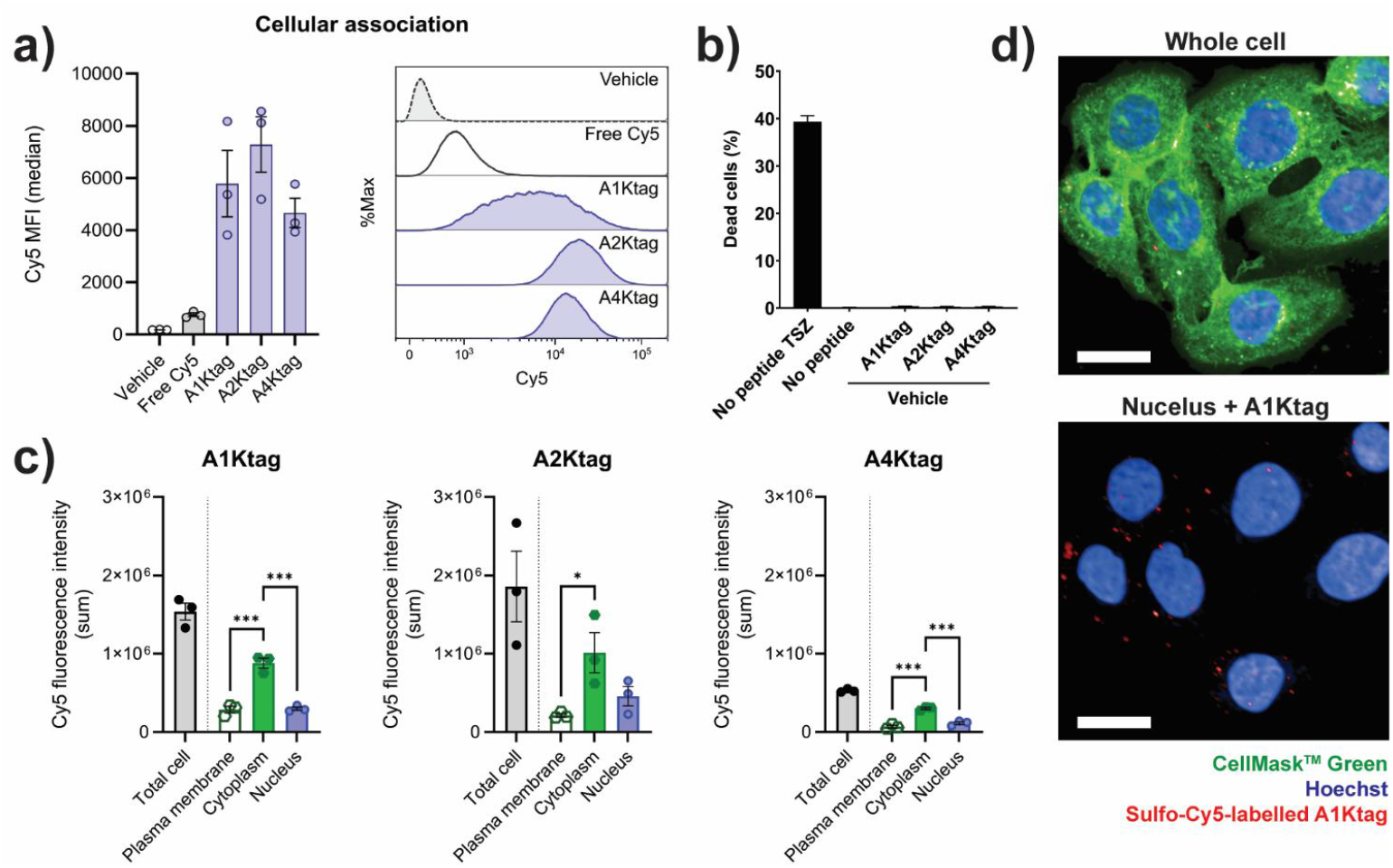
Results from incubating sulfo-Cy5-labelled tetra-lysine-tagged peptides with live HT-29 cells. a) Cellular association of 20 nM sulfo-Cy5-labelled peptides by flow cytometry showing median Cy5 intensity (left) and histogram of percent maximum Cy5 intensity (right). b) Percentage cell death detected by 2D imaging of propidium iodide staining following an 8-hour incubation of 50 nM sulfo-Cy5 labelled peptides with live HT-29 cells compared with cells treated for necroptosis. c) Analysis of peptide location within cells calculated from 3D imaging following a 4-hour incubation with live HT-29 cells. d) Maximum intensity projection of an example image used for analysis in (c) showing 20 nM Cy5-labelled A1Ktag peptides (red) and Hoechst nuclear stain (blue) with (top) and without (bottom) CellMask™ Green plasma membrane stain (green). Images taken at 63X magnification, scale bar represents 20 μm.

## Conclusion

Self-assembling amyloidogenic proteins are challenging targets, because many are intrinsically disordered as monomers, hence identification of molecules that bind to them through enthalpy-driven mechanisms is difficult. The end-point fibrils have a stable, ordered cross-β core but may be surrounded by flexible N- or C-termini. Along the course of fibril assembly, a heterogeneous mixture of oligomeric species is populated. In spite of these obstacles, cyclic peptides present promising amyloid-binding candidates because they have a constrained structure and will therefore experience a relatively small entropic loss upon binding to flexible regions, while offering the potential for many interactions with the extensive fibril surface. Mirror-image phage display and phage-assisted continuous evolution (PACE) have previously identified cyclic peptides that bind and inhibit formation of disease-associated amyloids^[28-30]^. Here we have successfully applied RaPID mRNA display to identify the first cyclic peptide binders for a functional amyloid, hRIPK3, during the process of transitioning from monomer to fibril. These peptides did not inhibit amyloid assembly but could bind to both monomeric and fibrillar forms of hRIPK3, as shown in the cell-based experiments, where peptide binding to hRIPK3 was observed under vehicle-treated, apoptosis and necroptosis conditions. The non-cytotoxic and cell permeable nature of the peptides allows them to be used to monitor hRIPK3 in cells under normal physiological conditions and to track hRIPK3 during programmed cell death. These peptides represent a useful starting point for the development of tools that can report on localisation and intracellular partners of hRIPK3 and be adapted to modulate its activity for therapeutic outcomes.

## Supporting Information

The authors have cited additional references within the Supporting Information^[31]^.

## Acknowledgements

This study was funded by NHMRC Ideas Grant (APP2019231) awarded to MS, MS and NS, and the Australian Research Council Centre of Excellence for Innovations in Peptide and Protein Science (project ID: CE200100012) awarded to R. J. P. JAB and OL were supported by Research Training Program stipends from the Australian Department of Education. The funders played no role in study design, data collection, analysis and interpretation of data, or the writing of this manuscript. The authors acknowledge the facilities and the expert scientific and technical assistance of staff within the Sydney Analytical, Sydney Microscopy & Microanalysis and Sydney Cytometry Core Research Facilities at the University of Sydney. We acknowledge and pay respect to the Gadigal people of the Eora Nation, the traditional owners of the land on which we research and collaborate at the University of Sydney.

