## Supporting Information for "*De novo* Cyclic Peptides Allow Visualisation of the Monomeric and Functional Amyloid Conformations of the Kinase RIPK3"

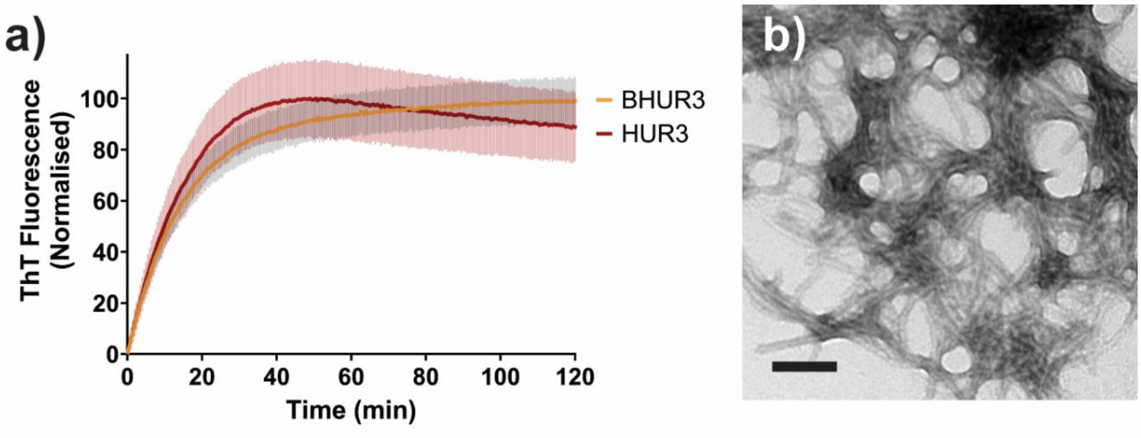

**Supplementary Figure 1:** ThT and TEM of BHUR3. a) ThT fluorescence from samples of 5 µM HUR3 or BHUR3 in PBS pH 7.4 at 37 ºC, normalised to plateau intensity. Error bars show SD. b) TEM of 0.2 mg/mL BHUR3 fibrils assembled in TBS, pH 8.0. Image taken at 110 000X magnification, scale bar represents 100 nm. Grid stained with 2% uranyl acetate.

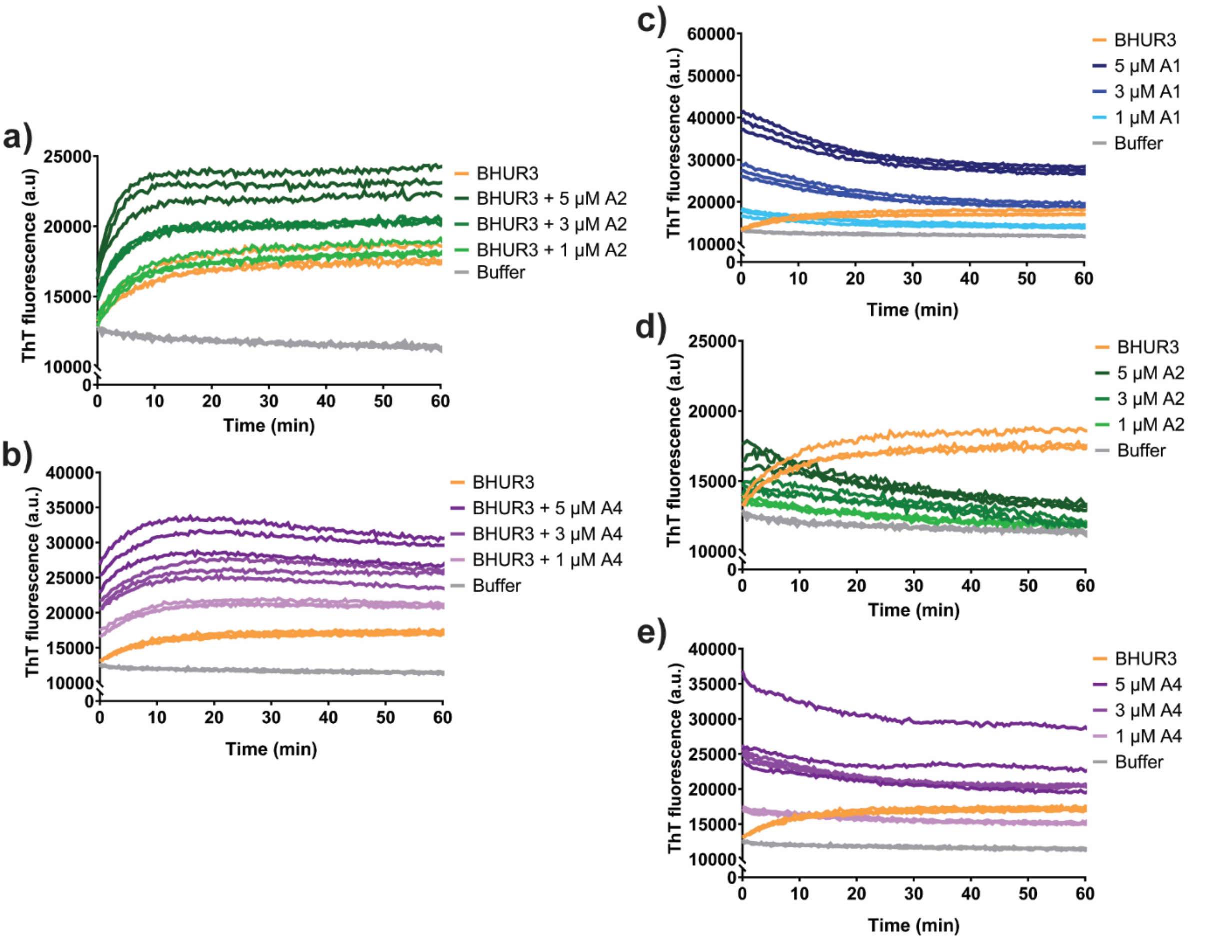

**Supplementary Figure 2:** ThT traces of RIPK3-binding peptides with and without BHUR3**.** Triplicate ThT traces of 1 μM BHUR3 co-incubated with 1, 3 or 5 μM a) A2, or b) A4. Triplicate ThT traces of 1 μM BHUR3 alone compared with 1, 3 or 5 μM c) A1, d) A2 or e) A4 alone. Assays performed at room temperature in PBS pH 7.4 supplemented with 40 μM ThT.

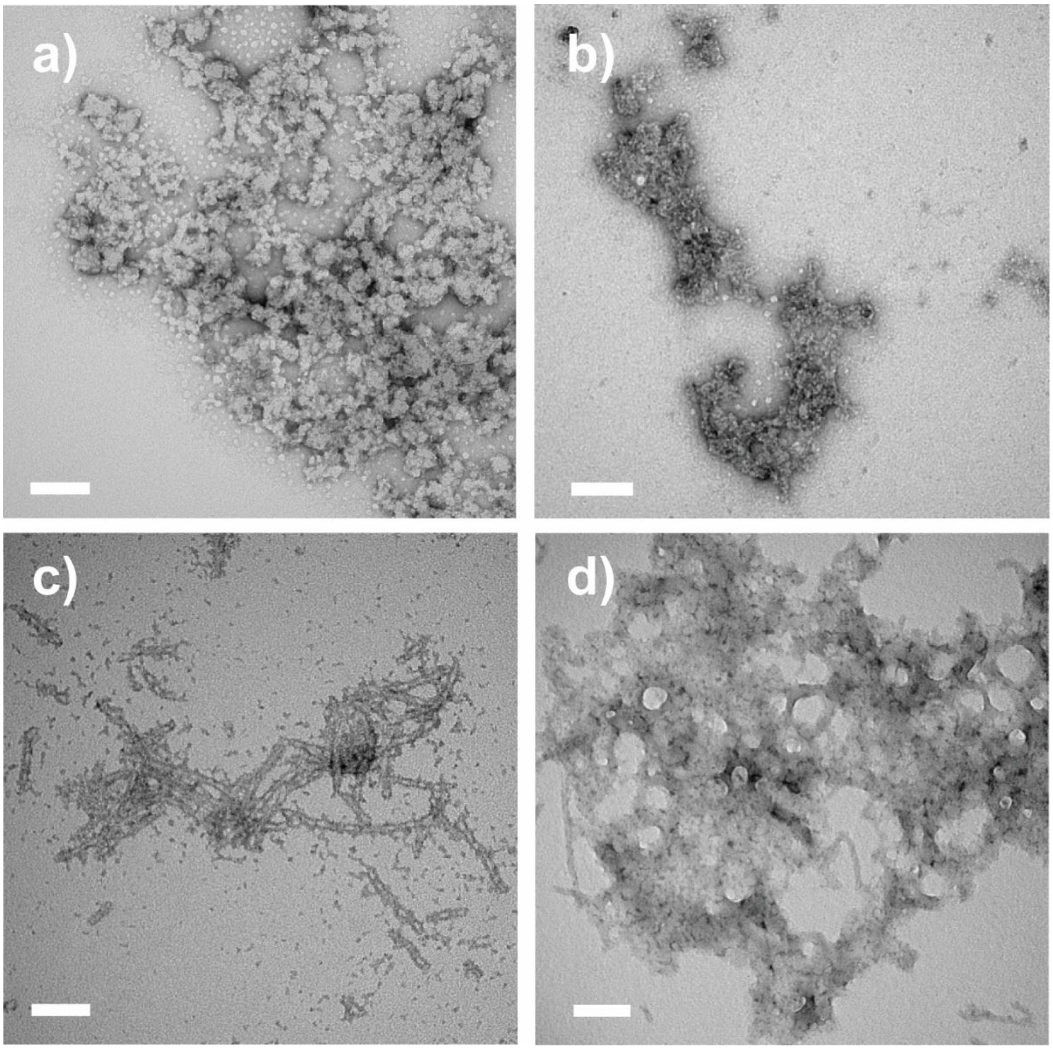

**Supplementary Figure 3:** TEM of A2 and A4 self-assembly with and without hRIPK3$_{387\text{-}518}$**.** TEM images of structures formed by a) 10 µM A2, and b) 10 µM A4 peptide alone. TEM images of hRIPK3$_{387–518}$ fibrils incubated with c) 2 µM A2, and d) 2 µM A4 peptide solutions. Images taken at 110 000X magnification, scale bars represent 100 nm. Grids were stained using 2% uranyl acetate.

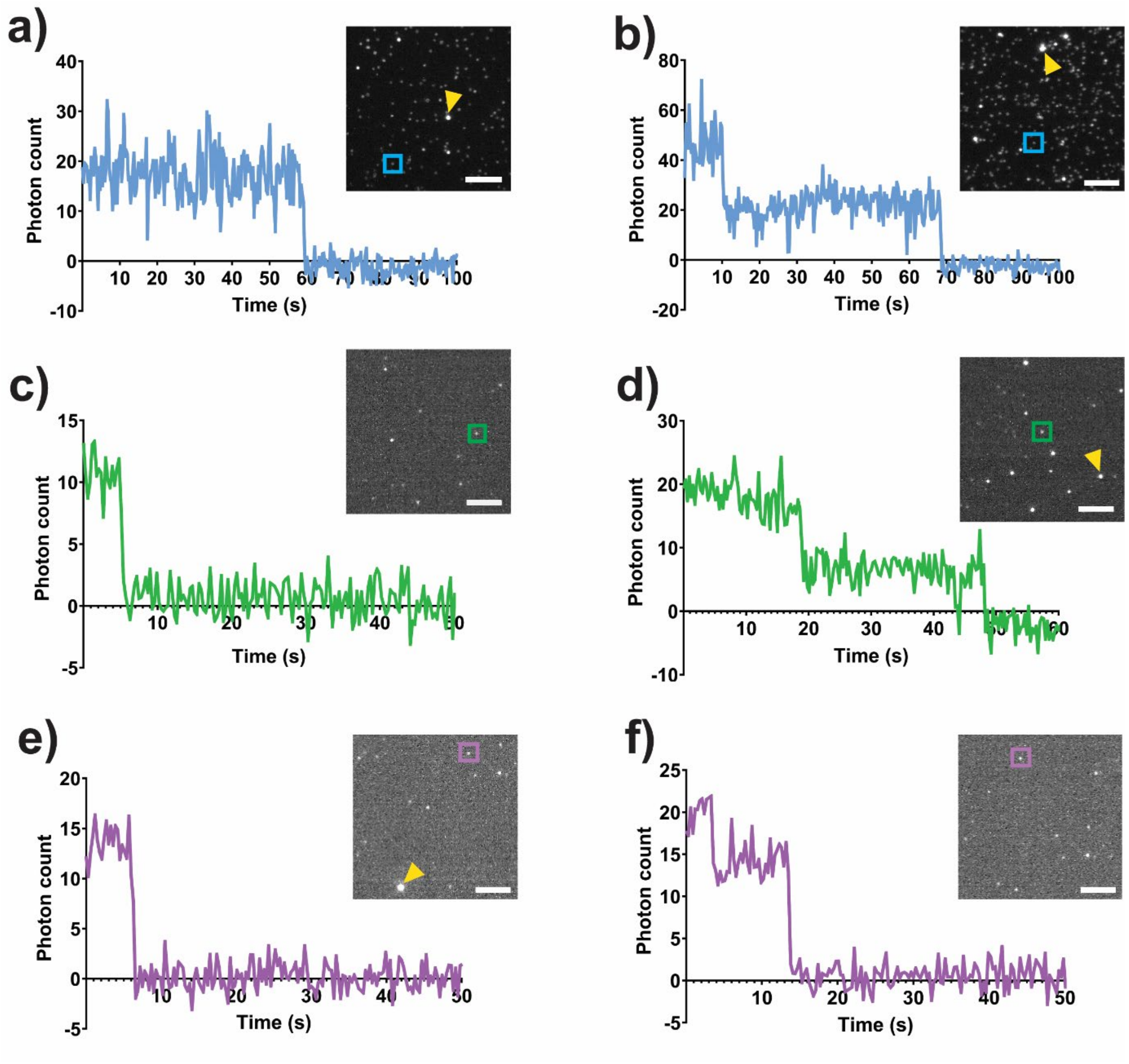

**Supplementary Figure 4:** Single molecule photobleaching of A1*, A2* and A4* peptides. Sulfo-Cy5-labelled A1* (50 pM) displaying a) monomeric, and b) dimeric species. Sulfo-Cy5-labelled A2* (50 pM) displaying c) monomeric, d) dimeric, and. Sulfo-Cy5-labelled A4* (100 pM) displaying e) monomeric, and f) dimeric. Coloured boxes indicate fluorescent spots corresponding to analysis shown. A1 in 50 mM Tris pH 8.0, 10 mM NaCl, 10% (w/v) D-glucose containing 1% GLOX solution (56 mg/mL glucose oxidase, 17 mg/mL catalyse, 10 mM Tris pH 8.0, 50 mM NaCl) and 10 mM MEA. A2 and A4 imaged without GLOX solution. Imaging performed in TIRF with 640 nm laser line (6% power), exposure at 300 ms/frame. Images corrected for background. Analysis performed using FIJI image analysis software. Images taken at 100X magnification, scale bars represent 5 μm. Examples of multimeric species with stoichiometry >4 indicated with yellow triangles.

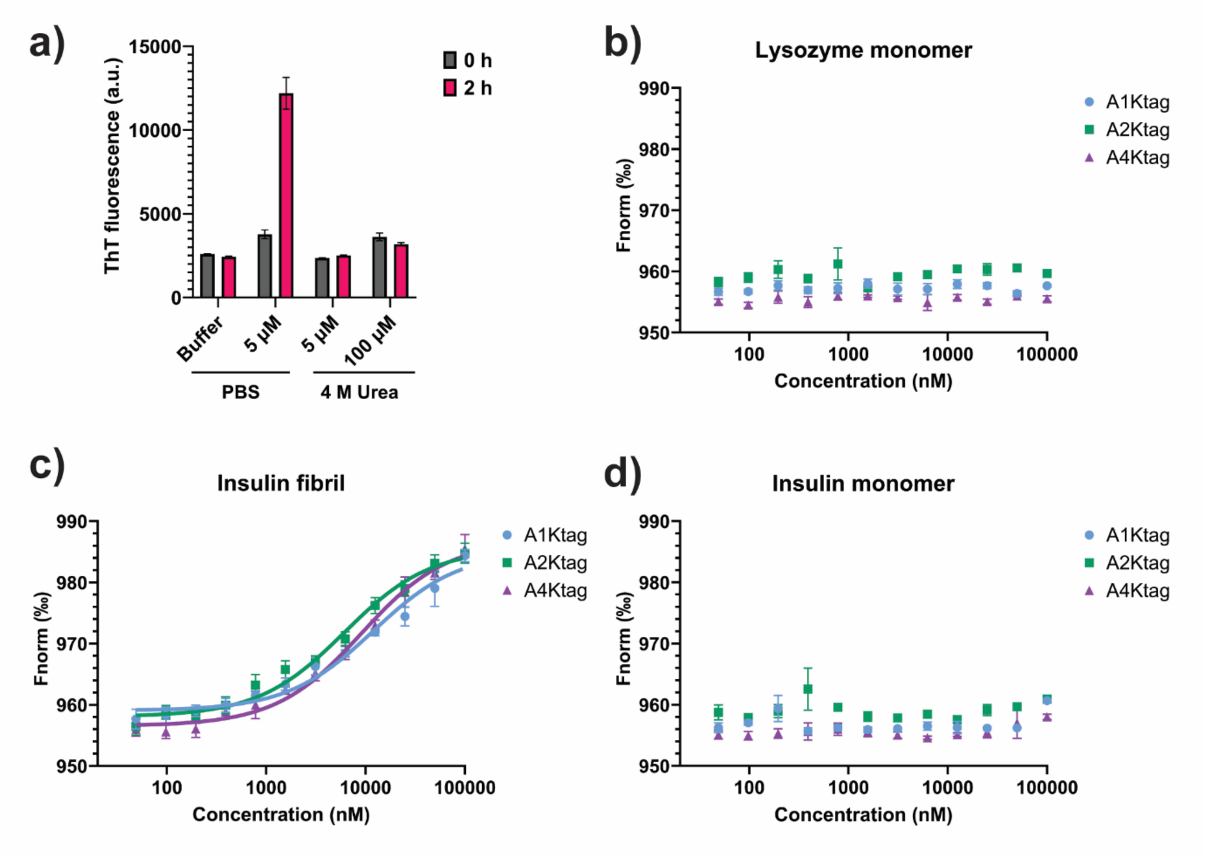

**Supplementary Figure 5:** MST data of control amyloid proteins**.** a) Mean initial (0 h) ThT fluorescence and endpoint fluorescence demonstrating monomeric nature of 5 µM and 100 µM HUR3 in 4 M urea over the course of 2 hours versus 5 µM HUR3 diluted into PBS. b-d) MST traces using 20 nM sulfo-Cy5-labelled tetra-lysine-tagged peptides against b) lysozyme monomer, c) insulin fibrils or d) insulin monomer. Raw data values shown as points with SD, fit values represented with solid line. Assays performed in triplicate, b-c) n = 2, d) n = 1. MST Assays were performed in TBS-T pH 8.0, 0.1 mg/mL BSA, 0.04% glycerol, 5% DMSO.

**Supplementary Table 1:** Summary of all calculated $K_d$ values from MST experiments

| Target | Peptide | $K_d$ (μM) | $K_d$ confidence (μM) |
|---|---|---|---|
| RIPK3 monomer | A1Ktag | 35.87 | ± 22.90 |
| | A2Ktag | 45.52 | ± 19.25 |
| | A4Ktag | 53.43 | ± 27.04 |
| IAPP fibril | A1Ktag | 49.02 | ± 9.77 |
| | A2Ktag | 25.26 | ± 3.51 |
| | A4Ktag | 24.80 | ± 4.80 |
| Insulin fibril | A1Ktag | 11.97 | ± 2.95 |
| | A2Ktag | 6.10 | ± 1.14 |
| | A4Ktag | 9.11 | ± 1.38 |

### Experimental methods

#### Production of Biotinylated (BHUR3) and Non-Biotinylated (HUR3) RHIM Domain of Human RIPK3

The RHIM-containing domain of hRIPK3 (hRIPK3$_{387\text{-}518}$) was expressed as a His$_6$-ubiquitin-TEV-RIPK3$_{387\text{-}518}$ construct (HUR3) or biotinylated His$_6$-ubiquitin-TEV-RIPK3$_{387\text{-}518}$ (BHUR3) in BL21(DE3) pLysS *E. coli* (Novagen), with cells grown at 37 °C for 3 h after induction with 0.5 mM IPTG at mid log phase. For complete biotinylation, 100 µM biotin was added during IPTG induction of BHUR3. BHUR3 and HUR3 were purified by Ni-NTA affinity chromatography under denaturing conditions. Expression pellets were resuspended in 20 mM Tris pH 8.0, 150 mM NaCl, 1 mM EDTA and lysed by sonication. Lysate was centrifuged at 26 000 × *g* for 45 min at 4 °C and insoluble material retained and resuspended in 6 M GuHCl, 20 mM Tris pH 8.0 with harsh agitation at room temperature (RT) until fully dissolved. Guanidine solution was centrifuged again at 26 000 × *g* for 30 min at 4 °C then, 1 mM β-mercaptoethanol was added and supernatant taken forward for Ni-NTA affinity chromatography. B(HUR3) was buffer exchanged into 8 M urea, 20 mM Tris, 100 mM $NaH_2PO_4$ occurred on the column and BHUR was eluted from the affinity resin by reduction of buffer to pH 4.0. Protein was concentrated to >200 µM using 10 kDa MWCO spin filters (Millipore), then buffer exchanged into 8 M urea 20 mM sodium acetate pH 4.0 and stored at 4 °C to avoid unwanted self-assembly. Concentration of BHUR3 and HUR3 was determined by $A_{280}$ measurement (using extinction coefficients of 34 950 $M^{-1}.cm^{-1}$ for BHUR3, 29 450 $M^{-1}.cm^{-1}$ for HUR3. Reagents were purchased from Sigma Aldrich unless otherwise indicated.

#### BHUR3 Fibril Formation and Removal of His$_6$-Ubiquitin

(Biotin)-His$_6$-ubiquitin-TEV-RIPK3$_{387\text{-}518}$ was diluted into 8 M urea, 20 mM sodium acetate pH 4.0 to ≤100 µM protein concentration and dialysed against 25 mM $NaH_2PO_4$, 150 mM NaCl (SnakeSkin dialysis tubing 3.5 kDa MWCO) containing 0.5 mM DTT for 1 h at RT, then Tobacco Etch Virus (TEV) enzyme (produced in house) was added at minimum of 75 µg/mL final concentration and samples were dialysed overnight with gentle magnetic stirring. Samples from before and after dialysis were analysed by SDS PAGE to assess extent of TEV cleavage. If incomplete, sample was resolubilised and this process was repeated. Fibrils were recovered by centrifugation at 17 000 × *g* for 10 min and washed into desired buffer twice to remove TEV and (Biotin-)His$_6$-ubiquitin tag, then stored for use at 4 °C. Cleaved BHU product was retained and concentration estimated for use in RaPID counterselection using $\varepsilon$ = 8480 $M^{-1}.cm^{-1}$.

#### RaPID Screening

Optimal bead loading concentrations were determined by incubating 2 pmol BHUR3 protein with 8, 4, 2, 1, 0.5 and 0.25 µL Streptavidin M280 DynaBeads™ and analysing immobilisation efficiency by SDS-PAGE with SYPRO™ Ruby protein stain. Optimal loading was determined by identifying the smallest volume of bead where no protein was observed in the supernatant fraction, indicating that all available protein had been immobilised.

For RaPID screening, a randomised cyclic peptide library was provided by Sydney Analytical Facilities (The University of Sydney). This library was prepared using *in vitro* transcription and translation from DNA oligos containing a fixed T7 promotor, a ribosome binding site and a start codon (AUG) then a random series of 4–15 NNK codons (where N = A, T, C or G and K = C or G), followed by a fixed cysteine-encoding codon (TGC) and region encoding a GNLI linker sequence. Briefly, *in vitro* transcription using T7 RNA polymerase was performed, and the resulting mRNA library was purified by denaturing urea polyacrylamide gel electrophoresis. mRNA library was ligated to a puromycin-linked oligonucleotide via T4 RNA ligase. mRNA-puromycin library (1 µM) was then translated to generate the cyclic peptide library via ribosomal synthesis using the PURExpress ΔRF kit (NEB), lacking RF1 in a Met-deficient system. This library was prepared using 25 µM initiating tRNA that had undergone aminoacylation with *N*-chloroacetyl L-tyrosine using eFx flexizyme such that all starting residues were translated as *N*-chloroacetyl L-tyrosine to allow for thioester formation with downstream cysteine residues[24, 31]. After 30 min at 37 °C, ribosomes were denatured by addition of 20 mM EDTA and peptides released into solution over another 30 min at 37 °C. mRNA-linked cyclic peptides subsequently underwent reverse transcription with M-MLV RNase (H–) reverse transcriptase (Promega) using appropriate reverse primer, then TBST selection buffer (20 mM Tris. HCl pH 8.0, 150 mM NaCl) containing 0.5 mM DTT was added so that final library volume was 300 µL. A small sample of this library (input) was reserved and diluted 100-fold for RT-qPCR analysis.

For round 1 of RaPID screening, BHUR3 was diluted out of denaturant into starting cyclic peptide library to a final concentration of 0.8 µM. This mixture was gently rotated at RT for 20 min, then overlayed onto Streptavidin M280 DynaBeads™ that had been prewashed thrice with selection buffer (0.25 µL bead slurry/pmol protein). This mixture was rotated at RT for a further 5–10 min to pull down biotinylated protein and bound peptides, then supernatant was removed. Beads were washed thrice in an equal volume of selection buffer changing tubes each wash, then resuspended in 100 µL 0.1% Triton X-100 and boiled at 95 °C for 5 min to recover bound peptides. A small sample (1 µL) peptide recovery mixture was reserved for RT-qPCR analysis.

For all rounds after the first, counterselection was performed prior to positive selection using TEV-cleaved BHU protein. 6X bead volume required for screening was washed thrice with selection buffer then incubated with 0.8 µM BHU for 30 min at RT gently rotating, then washed twice with selection buffer again. Beads were split into 3 aliquots (50 µL), and each was incubated with the prepared cyclic peptide library for 30 min gently rotating at RT to remove non-specific binders. Final bead aliquots were washed and recovered for RT-qPCR analysis as previously described. Positive selection for all subsequent rounds of screening occurred as for round 1 but in 50 µL total volume.

Recovered DNA was amplified by PCR with appropriate primers, and once sufficient DNA product was visualised, the PCR reaction products were precipitated out with 0.1 vol 3 M NaCl and 2.2 vol 100% ethanol. Precipitated DNA was recovered by centrifugation at 12 000 × g for 10 min, then washed with 70% (v/v) ethanol Pellet was allowed to air dry then resuspended in MilliQ water (10 µL). Resuspended DNA was used for transcription, with reactions containing reserved DNA (5 µL), 1X T7 buffer, DTT (10 mM), $MgCl_2$ (30 mM), (5 mM ea.) and 1X T7 RNA polymerase. Reaction mixtures were prepared on ice then incubated at 37 °C overnight. The reaction was quenched by the addition of NaCl (300 mM) and EDTA (25 mM), then mRNA was precipitated by addition of ⅞ volume 2-isopropanol and recovered by centrifugation at 15 000 × g for 5 min. Pellet was washed with 70% ethanol then allowed to air dry. mRNA pellet was resuspended in MilliQ water (20 µL) and concentration was determined by $A_{260}$ signal (average mRNA length 106 nt, molecular weight 340 Da). mRNA was diluted to 10 µM in MilliQ water and puromycin was ligated to transcribed mRNA. Puromycin ligation reactions contained 1X T4 buffer (2.1.1), prepared mRNA library (1 µM) and 0.15 U/µL T4 RNA ligase. Ligations occurred during incubation at 37 °C for 30 min then were quenched and precipitated by the addition of 300 mM NaCl, 25 mM EDTA and 250 µg/mL glycogen followed by 4 vol of 2-isopropanol. Precipitated mRNA puromycin conjugates were pelleted by 15 min centrifugation at 15 000 × g, washed once with 70% ethanol, then air dried. The pellet was resuspended in 2 µL MilliQ water to yield a stock concentration of ~6 µM. Translation of mRNA-puromycin libraries for all subsequent rounds was performed as for round 1 but at a 1/20 scale. Following translation, reverse transcription was completed by adding 250 µM ea. dNTPs, 15 µM $Mg(OAc)_2$, 25 mM Tris pH 8.0, 10 mM KOH, 2 µM reverse primer and 1X M-MLV RT (H–). Reaction was allowed to incubate at 37 °C for 30 min, then the cyclic peptide library was diluted to 50 µL in TBST buffer containing 0.5 mM DTT.

Positive recovery samples for all rounds were prepared and sent for sequencing by iSeq at the Ramaciotti Centre for Genomics at the University of New South Wales.

**General Materials and Methods for Peptide Synthesis**

Peptide grade *N,N*-dimethylformamide (DMF) for peptide synthesis was purchased from RCI. Gradient grade acetonitrile (MeCN) for chromatography was purchased from Sigma Aldrich and ultrapure water (Type 1) was obtained from a Merck Millipore Direct-Q 5 water purification system. Standard fluorenylmethoxycarbonyl (Fmoc)-protected amino acids (Fmoc-Xaa-OH), coupling reagents and resins were purchased from Mimotopes or Novabiochem. Fmoc-$PEG_2$-OH and Fmoc-Cha-OH were purchased from GL Biochem (Shanghai) Ltd and AK Scientific, respectively. Piperidine and *N,N*-di*iso*propylethylamine (DIPEA) were purchased from Alfa Aesar and Merck, respectively. Fmoc-SPPS was performed through automated synthesis on a Syro I peptide synthesizer (Biotage). Sulfo-Cy5 DBCO was purchase from BroadPharm. All other reagents were purchased from Sigma Aldrich, AK Scientific or Merck and used as received.

**Preparative Liquid Chromatography**

Preparative and semi-preparative reversed-phase high performance liquid chromatography (RP-HPLC) was performed using a Waters 600E multisolvent delivery system with a Rheodyne 7725i injection valve (5 mL loading loop) with a Waters 500 pump and a Waters 490E programmable wavelength detector operating at 214 nm and 280 nm. Preparative reversed-phase HPLC was performed using a Waters X-Bridge® C18 OBD™ Prep Column (5 µm, 30 × 150 mm) at a flow rate of 38 mL $min^{-1}$ using a mobile phase of 0.1% TFA in water (solvent A) and 0.1% TFA in MeCN (solvent B) on linear gradients, unless otherwise specified.

**Liquid Chromatography-Mass Spectrometry**

Liquid Chromatography-Mass Spectrometry (LC-MS) was performed on a Shimadzu 2020 UPLC-MS instrument with a Nexera X2 LC-30AD pump, Nexera X2 SPD-M30A UV/Vis diode array detector and a Shimadzu 2020 (ESI) mass spectrometer operating in positive ion mode. Separations were performed on a Waters Acquity BEH300 1.7 µm, 2.1 × 50 mm (C18) column at a flow rate of 0.6 mL $min^{-1}$. All separations were performed using a mobile phase of 0.1 vol.% formic acid in water (solvent A) and 0.1 vol.% formic acid in MeCN (solvent B) using linear gradients over 5 min.

**Analytical RP-HPLC**

Analytical reversed-phase HPLC was performed on a Waters Alliance e2695 HPLC system equipped with a 2998 PDA detector (λ = 210–400 nm). Separations were performed on a Waters XBridge® Peptide BEH300 5 µm, 4.6 × 250 mm (C18) column at 40 °C with a flow rate of 1.0 mL $min^{-1}$. All separations were performed using a mobile phase of 0.1% TFA in water (Solvent A) and 0.1% TFA in MeCN (Solvent B) using linear gradients, unless otherwise specified. Analytical RP-HPLC traces were processed where time 0 min refers to the start of the gradient.

**Mass Spectrometry**

Low resolution mass spectra were recorded on a Shimadzu 2020 (ESI) mass spectrometer operating in positive and negative mode. High resolution mass spectra were recorded on a Bruker-Daltronics Apex Ultra 7.0 T Fourier transform (FTICR) mass spectrometer.

**Peptide Synthesis**

**General procedure A;** Automated Peptide Synthesis (SYRO I peptide synthesizer): The resin (90 mg, 50 µmol, 0.56 mmol $g^{-1}$, 1 eq.) was treated with 40 vol.% piperidine in DMF (800 µL) for 4 min, drained, then treated with 20 vol.% piperidine in DMF (800 µL) for 4 min, drained, and washed with DMF (4 × 1.2 mL). The resin was then treated with a solution of Fmoc-Xaa-OH or chloroacetic acid (200 µmol, 4 eq.) and Oxyma (220 µmol, 4.4 eq.) in DMF (400 µL), a 1 wt.% solution of 1,3-diisopropyl-2-thiourea in DMF (400 µL), followed by a solution of DIC (200 µmol, 4 eq.) in DMF (400 µL). Coupling of Fmoc-Cys(Trt)-

OH and Fmoc-His(Trt)-OH were carried out at 50 °C for 30 min. All other coupling reactions were conducted at 75 °C for 15 min. The resin was then drained and washed with DMF (4 × 1.2 mL) before being treated with a solution of 5 vol.% $Ac_2O$ and 10 vol.% *i*$Pr_2$NEt in DMF (800 µL) for 6 min at room temperature, drained and washed with DMF (4 × 1.2 mL).

**General procedure B**: Manual cleavage: The resin was thoroughly washed with $CH_2Cl_2$ (5 × 5 mL) before being treated with 87.5:5:5:2.5 v/v/v TFA:tri*iso*propylsilane:$H_2O$:EDT (5 mL) and shaken at room temperature for 2 h. The resin was filtered, and the filtrate concentrated under a stream of nitrogen before addition of diethyl ether (40 mL). The peptide was pelleted by centrifugation (4 min, 4 °C, at 5000 × *g*) and the ether was decanted. The crude peptide was dissolved in the minimum volume of 1:1 MeCN/$H_2O$ and concentrated by lyophilisation.

**General procedure C:** Manual cleavage: The resin was thoroughly washed with $CH_2Cl_2$ (5 × 5 mL) before being treated with 90:5:5 v/v/v TFA:tri*iso*propylsilane:$H_2O$ (5 mL) and shaken at room temperature for 2 h. The resin was filtered, and the filtrate concentrated under a stream of nitrogen before addition of diethyl ether (40 mL). The peptide was pelleted by centrifugation (4 min, 4 °C, at 5000 × *g*) and the ether was decanted. The crude peptide was dissolved in the minimum volume of 1:1 MeCN/$H_2O$ and concentrated by lyophilisation.

**General procedure D:** Cyclisation: The crude peptide (25 µmol) was dissolved in DMSO (5 mL) and *i*$Pr_2$NEt (180 µL) was added. The peptide solution was heated in a water bath at 60 °C until the cyclisation was complete as judged by UPLC-MS analysis.

**General procedure E**: Acm removal: The purified peptide (1.2 µmol) was dissolved in a 1:1 v/v mixture of MeCN and $H_2O$ with 0.1 vol.% TFA (0.1 M) and AgOAc (40 eq., 48 µmol) was added. The mixture was shaken at room temperature until the reaction was complete as judged by UPLC-MS analysis. Dithiothreitol (60 eq., 72 µmol) was added to the reaction prior to RP-HPLC.

**General procedure F:** Strain promoted alkyne-azide cycloaddition (SPAAC) reaction: The peptide was added to one molar equivalent of DBCO-fluorophore in a 1:1 v/v mixture of DMSO and $H_2O$ (approximately 10 mg/mL). The mixture was stirred at room temperature until the reaction was complete as judged by UPLC-MS analysis.

**Thioflavin T Assays**

Solutions with appropriate buffering agent at pH 4.0–8.0 as specified, NaCl as required and containing 40 µM ThT were prepared in Corning™ black clear bottom 96-well plates (cat no. 3631). Peptides and proteins were diluted from concentrated stocks into buffer immediately prior to measurement. Final urea concentration and DMSO volume did not exceed 200 mM and 1% unless otherwise specified. Plates were sealed with optically clear sealing film (Corning™, cat no. 6575) and loaded into a BMG POLARstar Omega microplate reader maintained at RT or 37 °C. Plates were shaken for 30 s at 700 rpm before first read (unless otherwise specified), and wells were excited at 440 nm, with fluorescence emission measured at 480 nm (filters +/– 10 nm). The gain was set at 1–5% from a buffer and ThT-containing sample. Assays were continued until fluorescence emission plateaued.

**Negative Stain Transmission Electron Microscopy**

Carbon-coated copper 200-mesh grids coated with formvar support film (ProSciTech Pty Ltd.) were glow discharged at 25 mA for 30 s at desired charge. Grids were floated on 20 µL droplets containing 10 µM peptide or 0.2 mg/mL $hRIPK3_{387–518}$ for 2 min, washed briefly with filtered MilliQ water three times and then blotted and stained by floating on 2% aqueous uranyl acetate stain for 30–60 s and blotted dry. For pull down experiments, fibrils were incubated with grids and washed as described, then grids were further floated onto droplets containing 2 µM peptide, washed, then stained as described.

Transmission electron microscopy was performed on using FEI Tecnai T12 electron microscope operating at 120 kV at the Sydney Microscopy and Microanalysis Core Research Facility. Images were captured using a Veleta CCD camera and RADIUS 2.0 imaging software (EMSIS GmbH) or a CMOS camera with the DigitalMicrograph Software (GATAN).

**Circular Dichroism Spectroscopy**

Experiments were performed using a Jasco J 815 CD Spectropolarimeter and Spectra Manager$^{TM}$ software (Jasco). Samples contained a final concentration of 0.3 mg/mL protein, peptide or matched buffer in a 1-mm High Precision Cell quartz cuvette (Hellma Analytics). Spectra were acquired with 0.5 nm step resolution from 260–190 nm, at 20 nm/min, with sample maintained at 25 °C. Spectra were collected using only one accumulation but were acquired three times for each sample, apart from buffer alone samples, which were acquired only once. Data were analysed using Spectra AnalysisTM software (Jasco). Buffer-alone spectra were subtracted from peptide or protein spectra and data smoothed using the Savitzky-Golay method. The secondary structure content was analysed using the BestSel online predictive software.

**Insulin Fibril Formation**

Purchased, lyophilised human insulin (Sigma Aldrich, cat no. 91077C) was dissolved at 2.5–3 mg/mL in 20 mM glycine pH 2.0. This was stored at 4 °C and used as monomeric control or heated at 60 °C for 8 h with shaking at 800 rpm to convert to amyloid fibrils. Amyloid formation was confirmed by ThT assay.

**IAPP Fibril Formation**

Purchased, lyophilised human IAPP (Abcam, ab142398) was dissolved in deionised water and concentration was calculated using A280 signal and the extinction coefficient (ε) 1615 $M^{-1}cm^{-1}$, then diluted into 10 mM Tris pH 7.5 at 70 µM and allowed to form quiescently at RT over 16 h. Amyloid formation was confirmed by ThT assay and TEM.

**Microscale Thermophoresis**

HUR3 was cleaved by TEV enzyme and assembled into fibrils as described previously. The sample was then buffer exchanged into TBST by two rounds of centrifugation at 17 000 × g for 10 min, resuspending the fibril pellet in fresh TBST following each spin. Insulin monomer was prepared, and fibrils were formed as described previously. Insulin fibrils and monomeric protein were then diluted to 200 µM and dialysed overnight into TBS to avoid buffer mismatch. Lyophilized lysozyme (Sigma Aldrich) was dissolved in TBS, filtered with a 0.22 µm filter and diluted to appropriate starting concentration. IAPP fibrils and monomer were prepared as described previously. Fibrils were recovered and washed into TBST buffer via centrifugation at 21 000 × *g* for 30 min and concentrated to 200 µM for MST assays. IAPP monomer samples were doped with concentrated buffer solutions to achieve buffer matching and assays were performed immediately after thawing. RIPK3 and insulin fibril samples were probe sonicated immediately prior to serial dilutions with 2 × 20 s at 25% power, with 45 s rest on ice in between each sonication. IAPP fibrils were also sonicated immediately prior to serial dilutions at 15% power for 30 s total. Protein samples was serially diluted 1:1 into PCR tubes to make dilutions of 100–0.048 µM or 45–0.044 µM (IAPP monomer) in TBST containing 0.04% glycerol, 0.1 mg/mL BSA, and sulfo-Cy5 labelled peptide so that total volume was 5 µL and final peptide concentration was 20 nM. Samples were incubated for 5–10 min then loaded into premium capillaries (NanoTemper) for measurement.

Microscale thermophoresis (MST) was conducted with a NanoTemper Monolith. For MST measurements, excitation power (red LED laser) was set to 95%, and MST power (infrared laser) was set to 20% and lasted 15 s. Fluorescence readings were taken for an additional 5 s before and after switching on infrared laser. Each capillary was measured in triplicate, and the experiment was repeated once. Data was analysed using MO Affinity Analysis v2.3 utilising a $K_d$ fit model (NanoTemper).

**TIRF Fluorescence Imaging**

Cleaned glass coverslips (24 × 24 mm, 130–160 µm thickness, Epredia) were treated with poly-L lysine solution (0.01%) immediately prior to use. $RIPK3_{387-518}$ fibrils (≤ 5 µM) were incubated with poly-L lysine-treated coverslips for 15 min then coverslips were washed thrice with TBST and blotted to remove unbound material. Sulfo-Cy5-labelled peptides were diluted into TBST (50–500 pM) and incubated with coverslips in the dark for 20 min, then coverslips were washed thrice with TBST and blotted, then overlayed with 20 µL 50 mM Tris pH 8.0, 10 mM NaCl before mounting onto slides and sealing with epoxy resin for imaging.

TIRF imaging was performed using an ONI Nanoimager microscope using an incidence angle between 51–53o. Objective was overlayed with immersion oil (Immersol™ 518 F, ne = 1.518), and $RIPK3_{387–518}$ fibrils were imaged via their autofluorescent properties using 473 nm laser line at <10% power, sulfo-Cy5 labelled peptides were imaged using 640 nm laser line at <5% power. Exposure set to 100 ms. Image processing for TIRF images were performed using FIJI/ImageJ analysis software. Background removal and beam correction were performed via image subtraction of the Gaussian blur (radius Σ = 80) of the TIRF image from the original.

**Single Molecule Photobleaching**

Cleaned glass coverslips (24 × 24 mm, 130–160 µm thickness, Epredia) were submerged in 100% acetone (~50 mL) in coplin jars. 2 mL silane (A3648) was added, and coplin jars were gently agitated to mix solution for 5 min then bath sonicated for 1 min. Coverslips were left to incubate in silane/acetone solution for a further 10 min then gently rinsed in MilliQ water and dried using $N_2$ gas. Sulfo-Cy5-labelled A1*, A2* or A4* peptides (50–100 pM) were diluted out of DMSO into 50 mM Tris pH 8.0, 10 mM NaCl, then overlaid over coverslips which were mounted onto glass slides and sealed immediately with epoxy resin. Imaging was performed at TIRF angles using the 640 nm laser line at 6% power, with 100 ms exposure per frame until all detected fluorophores had bleached. Hyperstack images were analysed using the single molecule biophysics plugin on FIJI/ImageJ analysis software to identify molecule positions and measure photon count over time.

**Immunofluorescence Assays Using HT-29 Cell Models of Cell Death**

HT-29 cells grown in TC treated T175 flasks (Corning™) were aspirated, washed once with RT DPBS then trypsinised with 0.3 vol warmed trypsin EDTA for 3–5 min at 37 °C twice. Trypsin was quenched with 0.7 vol warmed D10 and then the mixture was centrifuged 400 × g at RT for 5 min. Supernatant was discarded, then cells were resuspended in warm supplemented DMEM. A small sample was diluted 1:10 in trypan blue and loaded onto a haemocytometer for cell counting, then the cell suspension was diluted in warmed DMEM to $8.5\times10^4$ cells/mL. Cells were seeded onto a 96-well clear bottom PhenoPlate (Revvity) coated in a collagen I matrix at $1.7\times10^4$ cells/well and incubated at 37 °C in 5% $CO_2$ overnight.

CellTracker™ Orange CMRA (Invitrogen) was diluted to 5 µM in DMEM and warmed to 37 °C. HT-29 cells were washed once in DPBS, then CellTracker™ Orange (50 µL) was added to each well and incubated at 37 °C in 5% $CO_2$ for 45 min. Following incubation, CellTracker™ Orange solution was aspirated and replaced with 150 µL D10. Cells were treated to induce no cell death (0.125% DMSO vehicle), apoptosis (30 ng/µL hTNF + 1 µM BV6) or necroptosis (30 ng/µL hTNF + 1 µM BV6 + 25 µM zVAD-fmk), and then incubated at 37 °C in 5% $CO_2$ for 6–7 h.

Treated and untreated cells were washed twice with 100 µL DPBS at RT. HT-29 cells were fixed with 4% PFA (100 µL) in the dark at RT for 25 min, then washed once with TBS-T before being permeabilised with ice cold absolute methanol (100 µL) in the dark at –30 °C for 10 min. Following fixation and permeabilization steps, cells were washed twice then stored in cold TBST, and plates were stored at 4 °C in protected from light overnight. Cells were blocked before the addition of peptide with 20% NDS solution diluted into TBST. 20% NDS solution (50 µL) was incubated with each well for 30 min at RT, then wells were washed twice in TBST before the addition of peptides.

Peptide master mixes were prepared in blocking buffer to 10 nM (10% final DMSO concentration) immediately prior to addition to wells. Peptide master mixes (50 µL) were added to each well and incubated in the dark at RT for 30 min, then washed twice with TBST. DAPI stain was prepared to 1 µg/mL in TBST and added (50 µL) added to each well. Plates were incubated in the dark at RT for 30 min, then were sealed with optically clear sealing film and stored at 4 °C until imaging.

**Imaging and Analysis of IFA**

Imaging was performed on an Opera Phenix (PerkinElmer) at 63X magnification (water immersion) in confocal mode using the parameters given below. Z-stacking was performed at 1 µm intervals spanning 3 µm total distance.

| Dye | Excitation (nm) | Emission filter (nm) | Laser power (%) | Exposure (ms) |
|---|---|---|---|---|
| CellTracker Orange | 561 | 570/50 | 100 | 100 |
| DAPI | 405 | 435/40 | 50 | 100 |
| Alexa Fluor 488 | 488 | 500/40 | 100 | 100 |
| Cy5 | 640 | 650/100 | 100 | 300 |

Image analysis was performed using Harmony 5.1 PhenoLOGIC analysis software (PerkinElmer). Intensity thresholds were adjusted as indicated to remove background fluorescence based on control wells for image exports. Analysis was performed on all whole wells. Cell counts were identified using DAPI stain (Method C) and cytoplasm identified by CellTracker™ Orange stain (Method B), and all subsequent analysis was conducted on Cy5 fluorescence detected within cells as determined by these parameters. Total Cy5 intensity analysis and well as spot analysis (spots detected using Method B) was performed measuring number of detected spots, mean spot size ($px^2$) and spot fluorescence intensity (corrected and relative). Control peptide wells were excluded from image analysis due to the lack of cytoplasmic marker for cell threshold detection. Statistical significance between untreated, apoptosis and necroptosis induced conditions and between no peptide controls was determined by 2-way ANOVA testing using Tukey's multiple comparisons tests.

**Flow Cytometry**

HT-29 cells grown in TC-treated T75 flasks (Corning) were aspirated, washed once with RT DPBS then trypsinised with 0.4 vol warmed TrypLE™ Express (Gibco) 3–5 min at RT. Trypsin was quenched with 0.8 vol cDMEM10 media then cell suspension was centrifuged for 5 min 300 × *g* at RT. Supernatant was discarded, then cells were resuspended in 0.8 vol cDMEM. Cell suspension was filtered through a 70 µm cell strainer then a small aliquot was diluted 1:9 in trypan blue and loaded onto a haemocytometer for cell counting. Cells were diluted into warmed cDMEM to $0.1\times10^6$ cells/mL, then 1 mL was seeded in duplicate for each condition (100 000 cells/well) onto a clear, TC-treated 12-well plate (Corning). Plate was incubated at 37 °C + 5% $CO_2$ for 18–24 h.

Sulfo-Cy5-labelled peptides, free sulfo-Cy5 control or DMSO vehicle were incubated for 30 min at 250 nM in warmed serum-free Opti-MEM media. Cells were washed once with warmed serum-free Opti-MEM media then peptide or control master mixes were added to a final concentration of 20 nM. Final DMSO concentration was 1%. Plates were incubated at 37 °C + 5% $CO_2$ for 4 h. After incubation, media was aspirated and cells were trypsinised with 0.1 vol warmed Trypsin-EDTA for 5 min at 37 °C. Trypsin was quenched with 0.1 vol warmed cDMEM10 media then cells were transferred to a 96-well V-bottom plate and centrifuged for 3 min at 350 × *g* at RT. Supernatant was removed and cells were resuspended in an equal volume of PBS and centrifuged again for 3 min at 350 × *g* at RT. A control sample was also prepared with a 1:1 suspension of untreated cells and cells that had been heated 56 °C for 10 min Supernatant was removed, and 50 µL LIVE/DEAD™ Fixable Blue Stain (Invitrogen) (1:400 dilution in PBS), was added to cells. Cells were incubated for 30 min at RT, protected from light, then 100 µL PBS was added and cells were centrifuged 3 min at 350 × *g* at RT. Supernatant was removed, and cells were fixed by incubation of 100 µL 4% formaldehyde for 15–20 min at RT, protected from light. Cells were washed twice with 100 µL PBS and centrifugation for 5 min 500 × *g* at RT. Cells were resuspended in 25 µL residual supernatant and transferred to 1.5 mL tubes. Wells were rinsed with 75 µL PBS to collect remaining cells, then samples stored on ice in the dark. Samples were taken for data acquisition on the LSR Fortessa (BD) flow cytometer, acquiring a minimum of 10 000 events for each sample.

**Confocal Microscopy of Fixed HT-29 Cells with Peptide Added to Live Cells**

Cells were prepared as described previously in preparation for flow cytometry. For confocal microscopy, cells were diluted to $5\times10^4$ cells/mL. 200 µL was seeded in triplicate for each condition (10 000 cells/well) onto a black, optically clear flat bottom 96-well PhenoPlate (Revvity). Additional DPBS was added to the outer wells of the plate to minimise evaporation. Peptide and controls were added following incubation period as described previously.

After 4-hour incubation with peptides or controls, cells were washed with 200 µL serum-free Opti-MEM media then stained with 100 µL CellMask™ Green plasma membrane stain (Invitrogen) (1:1000 dilution in Opti-MEM media) for 5–10 min at 37 °C. Cells were fixed in 100 µL warmed 4% formaldehyde at 37 °C for 5–10 min then washed thrice with 100 µL PBS. 50 µL Hoechst 33324 solution (Invitrogen) (1:400 dilution in PBS) was added to washed cells and incubated for 5 min at RT, then cells were washed thrice again in 100 µL PBS. Plates were stored at 4 °C protected from light and imaged within 24 h using an Opera Phenix (PerkinElmer) at 63X magnification (water immersion) in confocal mode with appropriate excitation/emission filters. Z-stacking was performed at 0.5 µm intervals spanning 7 µm total distance.

Images were used to calculate subcellular localisations peptides within cells at the plasma membrane, within the cytoplasm or within the nucleus using the Harmony 5.1 PhenoLOGIC analysis software (PerkinElmer). Nuclei were identified by Hoechst staining, boundaries of the plasma membrane were defined by CellMask™ Green staining, and the cytoplasm was classes as within the bounds between the nuclei and the cell membrane. Statistical significance between subcellular localisations for each peptide was determined by one-way ANOVA testing using Tukey's multiple comparisons tests.

**2D Live-Cell Imaging of HT-29 Cells Following Peptide Incubation and TSV Treatment**

HT-29 cells were prepared in a 96-well black, optically clear flat bottom plate (Revvity) as described previously for confocal microscopy experiments. Peptides were preincubated with Opti-MEM media for 15 min at RT before being added to cells in media to a final concentration of 50 nM. Cells were incubated with peptides at 37 °C + 5% $CO_2$ for 4 h. Following incubation, cells were washed once with 100 µL serum-free FluoroBrite™ DMEM media (FB-DMEM), then CellTracker™ Green CMFDA (1:400 dilution) and Hoechst 33342 (1:10 000 dilution) diluted in warmed FB-DMEM media was incubated with cells for 30 min at 37 °C + 5% $CO_2$. After incubation, staining solution was removed and replaced with 100 µL warmed cFB-DMEM5 media supplemented with ProLong Live Antifade reagent (1:1000 dilution). Cells were treated with warmed TSZ solution (25 ng/mL hTNF, 1 µM BV-6, 25 µM zVAD-fmk) or vehicle (DMSO) in cFB-DMEM5 media containing 5 µg/mL propidium iodide. Plates were sealed with Breath-Easy™ gas-permeable plate membrane (brand) then loaded into the Opera Phenix Plus (PerkinElmer) with environmental controls (37 °C + 5% $CO_2$). Cells were imaged immediately using the 10X air objective (NA 0.3) and parameters given below. Images were acquired of 3–9 FOVs with 3 × 7.4 μm z-planes per well at 10–30 min intervals for 10 h.

| Dye | Excitation (nm) | Emission (nm) | Laser power (%) | Exposure (ms) |
|---|---|---|---|---|
| Hoechst 33342 | 361 | 497 | 75 | 100 |
| CellTracker Green | 492 | 517 | 50 | 100 |
| Propidium iodine | 535 | 617 | 100 | 100 |
| Cy5 | 647 | 665 | 100 | 100 |

### Peptide Characterisation

#### Synthesis of RIPK3-Binding Cyclic Peptides

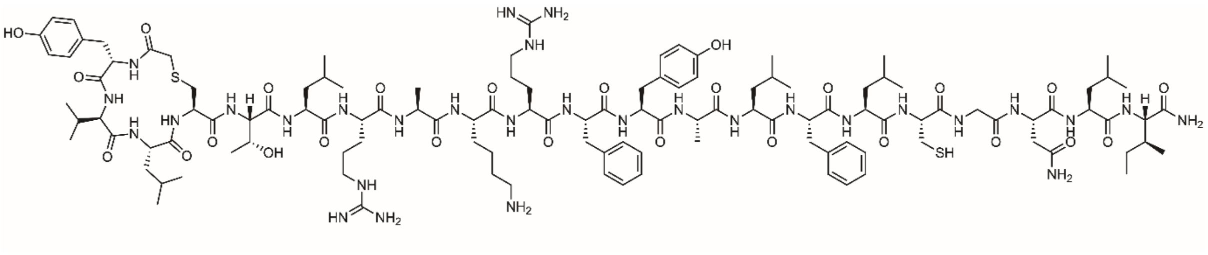

**A1:** Rink amide resin (75 mg, 50 µmol, 0.67 mmol $g^{-1}$) was loaded with Fmoc-Ile-OH according to general procedure **A**. The target peptide was then synthesised by iterative Fmoc-SPPS on the Syro I peptide synthesizer according to general procedure **A**, with chloroacetic acid as the final coupling step to generate the resin-bound sequence YVLCTLRAKRFYALFLC*GNLI where C* is Acm-protected Cys. The peptide was cleaved from resin by general procedure **B** and a portion (12.5 µmol) was cyclised according to general procedure **D**. Purification by preparative RP-HPLC (0 to 60 vol.% MeCN in $H_2O$ with 0.1 vol.% TFA over 50 min), followed by lyophilisation, afforded the pure peptide as a white fluffy solid (13.4 mg, 38%). A portion of this peptide (1.3 µmol) was taken forwards for Acm removal according to general procedure **E**. Purification by preparative RP-HPLC (0 to 75 vol.% MeCN in $H_2O$ with 0.1 vol.% TFA over 30 min), followed by lyophilisation, afforded the pure peptide as a white fluffy solid (0.7 mg, 21%). **LRMS:** (+ESI) *m/z* 1678.7 $[2M+3H]^{3+}$, 1259.3 $[M+2H]^{2+}$,1259.3 $[M+3H]^{3+}$. **Analytical RP-HPLC:** $R_t$ = 21.0 min (0 to 75 vol.% MeCN in $H_2O$ with 0.1 vol.% TFA over 30 min, λ = 214 nm).

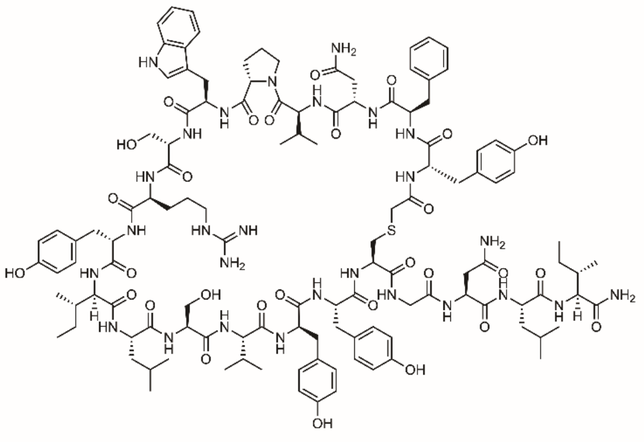

**A2:** Rink amide resin (75 mg, 50 μmol, 0.67 mmol $g^{-1}$) was loaded with Fmoc-Ile-OH according to general procedure **A**. The target peptide was then synthesised by iterative Fmoc-SPPS on the Syro I peptide synthesizer according to general procedure **A**, with chloroacetic acid as the final coupling step to generate the resin-bound sequence YFNVPWSRYILSVYYCGNLI. The peptide was cleaved from resin by general procedure **B** and a portion (12.5 μmol) was cyclised according to general procedure **D**. Purification by preparative RP-HPLC (0 to 65 vol.% MeCN in $H_2O$ with 0.1 vol.% TFA over 60 min), followed by lyophilisation, afforded the pure peptide as a white fluffy solid (6.2 mg, 19%). **LRMS:** (+ESI) *m/z* 1674.0 $[2M+3H]^{3+}$, 1255.7 $[M+2H]^{2+}$, 837.3 $[M+3H]^{3+}$. **Analytical RP-HPLC:** $R_t$ = 25.1 min (0 to 60 vol.% MeCN in $H_2O$ with 0.1 vol.% TFA over 30 min, λ = 214 nm).

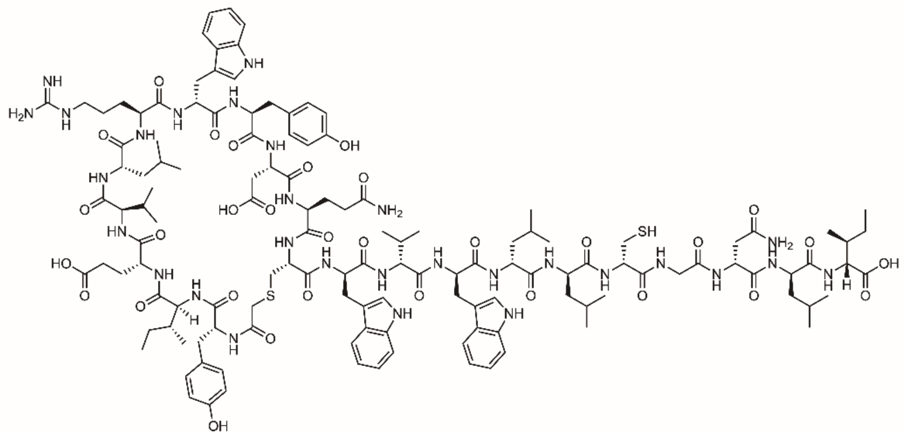

**A4:** Rink amide resin (75 mg, 50 µmol, 0.67 mmol $g^{-1}$) was loaded with Fmoc-Ile-OH according to general procedure **A**. The target peptide was then synthesised by iterative Fmoc-SPPS on the Syro I peptide synthesizer according to general procedure **A**, with chloroacetic acid as the final coupling step to generate the resin-bound sequence YIEVLRWYDQCWVWLLC*GNLI where C* is Acm-protected Cys. The peptide was cleaved from resin by general procedure **B** and a portion (12.5 µmol) was cyclised according to general procedure **D**. Purification by preparative RP-HPLC (0 to 65 vol.% MeCN in $H_2O$ with 0.1 vol.% TFA over 60 min), followed by lyophilisation, afforded the pure peptide as a white fluffy solid (3.4 mg, 38%). A portion of this peptide (1.2 µmol) was taken forwards for Acm removal according to general procedure **E**. Purification by preparative RP-HPLC (0 to 90 vol.% MeCN in $H_2O$ with 0.1 vol.% TFA over 90 min), followed by lyophilisation, afforded the pure peptide as a white fluffy solid (0.8 mg, 24%). **LRMS:** (+ESI) *m/z* 1817.5 $[2M+3H]^{3+}$, 1363.3 $[M+2H]^{2+}$, 909.1 $[M+3H]^{3+}$. **Analytical RP-HPLC:** $R_t$ = 23.3 min (0 to 75 vol.% MeCN in $H_2O$ with 0.1 vol.% TFA over 30 min, λ = 214 nm).

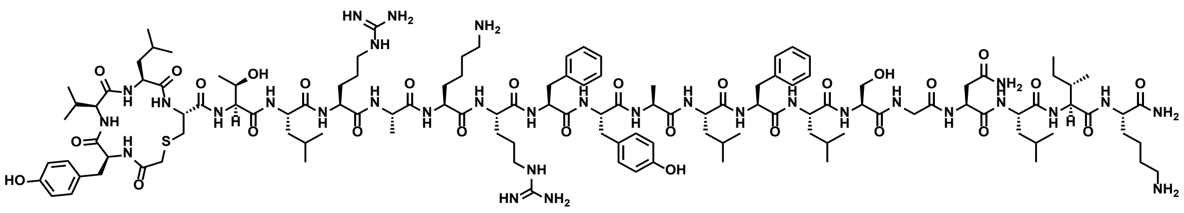

**A1*:** Rink amide resin (90 mg, 50 µmol, 0.56 mmol g$^{-1}$) was loaded with Fmoc-Lys(Boc)-OH according to general procedure **A**. The target peptide was then synthesised by iterative Fmoc-SPPS on the Syro I peptide synthesizer according to general procedure **A**, with chloroacetic acid as the final coupling step to generate the resin-bound sequence YVLCTLRAKRFYALFLSGNLIK. The peptide was cleaved from resin by general procedure **C** and a portion (25 µmol) was cyclised according to general procedure **D**. Purification by preparative RP-HPLC (0 to 50 vol.% MeCN in $H_2O$ with 0.1 vol.% TFA over 50 min), followed by lyophilisation, afforded the pure peptide as a white fluffy solid (17.9 mg, 23%). **HRMS:** Calculated for [$C_{125}H_{198}N_{32}O_{28}S$+H]$^+$: 2628.4847, found 2628.4782. **LRMS:** (+ESI) *m/z* 1753.9 [2M+3H]$^{3+}$, 1315.6 [M+2H]$^{2+}$, 877.3 [M+3H]$^{3+}$, 658.1 [M+4H]$^{4+}$. **Analytical RP-HPLC:** $R_t$ = 21.6 min (0 to 50 vol.% MeCN in $H_2O$ with 0.1 vol.% TFA over 30 min, λ = 214 nm).

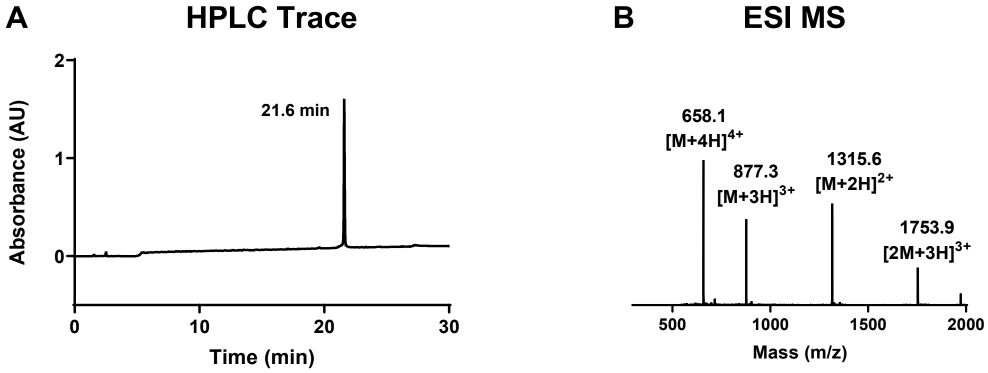

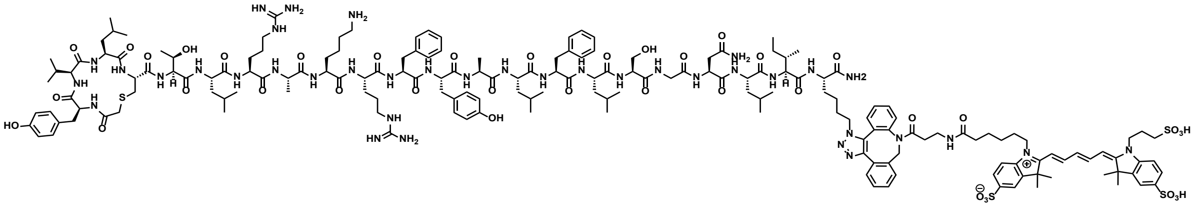

**Sulfo-Cy5-labelled A1*:** Rink amide resin (90 mg, 50 µmol, 0.56 mmol $g^{-1}$) was loaded with Fmoc-Lys*-OH according to general procedure **A** where Lys* refers to azidolysine. The target peptide was then synthesised by iterative Fmoc-SPPS on the Syro I peptide synthesizer according to general procedure **A**, with chloroacetic acid as the final coupling step to generate the resin-bound sequence YVLCTLRAKRFYALFLSGNLIK*. The peptide was cleaved from resin by general procedure **C** and a portion (25 µmol) was cyclised according to general procedure **D**. Purification by preparative RP-HPLC (0 to 50 vol.% MeCN in $H_2O$ with 0.1 vol.% TFA over 50 min), followed by lyophilisation, afforded the pure peptide as a white fluffy solid (6.6 mg, 9%). DBCO-Sulfo-Cy5 was attached to the peptide via a SPAAC reaction (procedure **F**), purified by preparative RP-HPLC (0 to 50 vol.% MeCN in $H_2O$ with 0.1 vol.% TFA over 50 min) and lyophilised, isolating the final peptide as a blue fluffy solid (3.8 mg, 96%). **HRMS:** Calculated for $[C_{177}H_{252}N_{38}O_{39}S_4+H]^+$: not found. **LRMS:** (+ESI) *m/z* 1833.3 $[M+2H]^{2+}$, 1222.5 $[M+3H]^{3+}$. **Analytical RP-HPLC:** $R_t$ = 21.4 min (0 to 70 vol.% MeCN in $H_2O$ with 0.1 vol.% TFA over 30 min, λ = 214 nm).

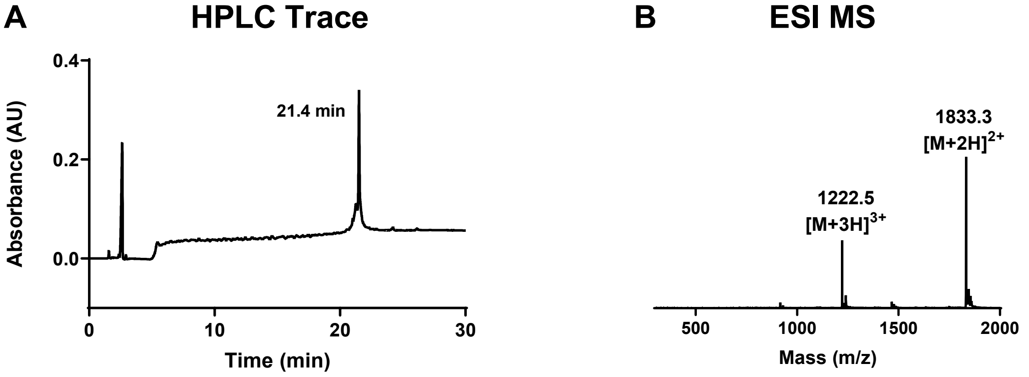

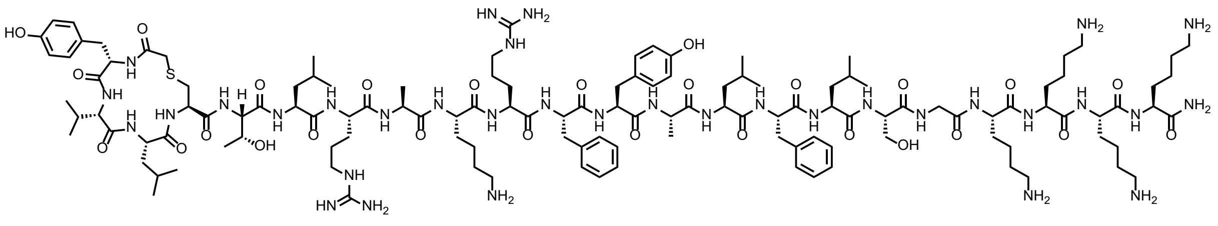

**A1Ktag:** Rink amide resin (90 mg, 50 µmol, 0.56 mmol $g^{-1}$) was loaded with Fmoc-Lys(Boc)-OH according to general procedure **A**. The target peptide was then synthesised by iterative Fmoc-SPPS on the Syro I peptide synthesizer according to general procedure **A**, with chloroacetic acid as the final coupling step to generate the resin-bound sequence YVLCTLRAKRFYALFLSGKKKK. The peptide was cleaved from resin by general procedure **C** and a portion (5 µmol) was cyclised according to general procedure **D**. Purification by preparative RP-HPLC (0 to 60 vol.% MeCN in $H_2O$ with 0.1 vol.% TFA over 50 min), followed by lyophilisation, afforded the pure peptide as a white fluffy solid (4.1 mg, 22%). **HRMS:** Calculated for $[C_{127}H_{206}N_{34}O_{27}S+H]^+$: 2672.5585, found 2672.5580. **LRMS:** (+ESI) *m/z* 1783.1 $[2M+3H]^{3+}$, 1337.4 $[M+2H]^{2+}$, 891.8 $[M+3H]^{3+}$, 668.9 $[M+4H]^{4+}$, 535.3 $[M+5H]^{5+}$, 446.3 $[M+6H]^{6+}$. **Analytical RP-HPLC:** $R_t$ = 16.8 min (0 to 70 vol.% MeCN in $H_2O$ with 0.1 vol.% TFA over 30 min, λ = 214 nm).

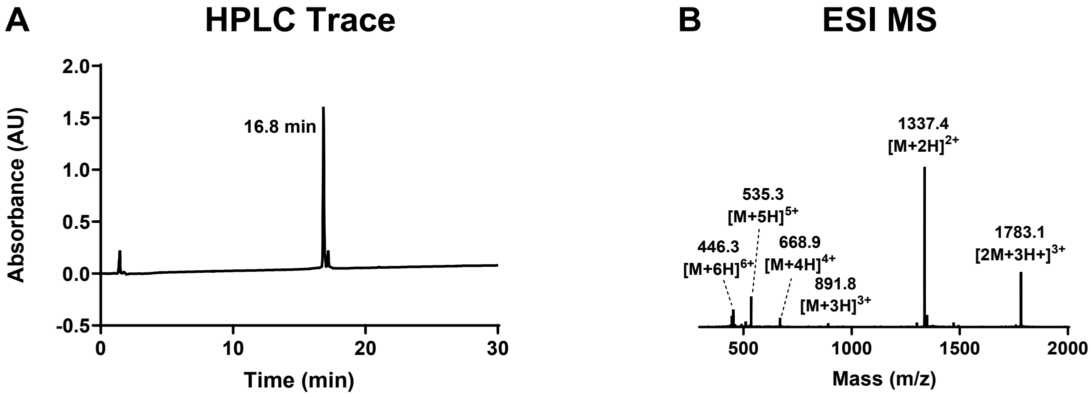

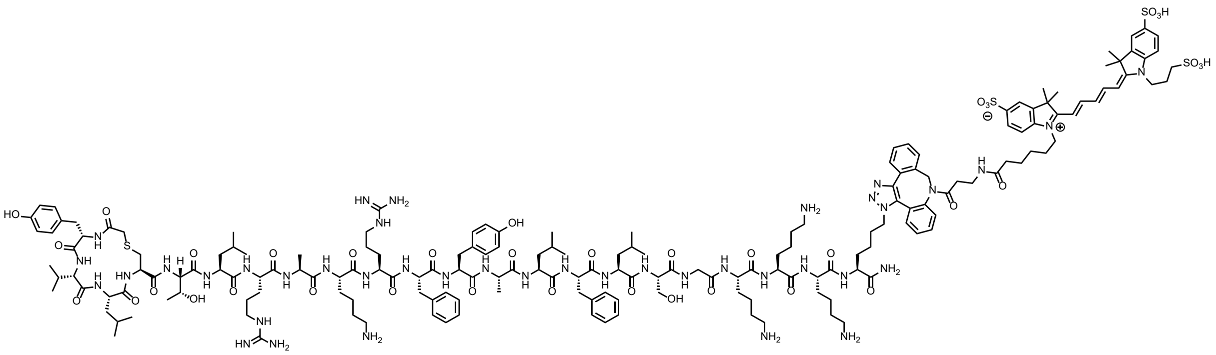

**Sulfo-Cy5-labelled A1Ktag:** Rink amide resin (90 mg, 50 µmol, 0.56 mmol g$^{-1}$) was loaded with Fmoc-Lys*-OH according to general procedure **A** where Lys* refers to azidolysine. The target peptide was then synthesised by iterative Fmoc-SPPS on the Syro I peptide synthesizer according to general procedure **A**, with chloroacetic acid as the final coupling step to generate the resin-bound sequence YVLCTLRAKRFYALFLSGKKKK*. The peptide was cleaved from resin by general procedure **C** and a portion (5 µmol) was cyclised according to general procedure **D**. Purification by preparative RP-HPLC (0 to 60 vol.% MeCN in $H_2O$ with 0.1 vol.% TFA over 50 min), followed by lyophilisation, afforded the pure peptide as a white fluffy solid (3.6 mg, 21%). DBCO-Sulfo-Cy5 was attached to the peptide via a SPAAC reaction (procedure **F**) and purified through a silica plug. Following lyophilisation, the final peptide was isolated as a blue fluffy solid (1.4 mg, 58%). **HRMS:** Calculated for $[C_{179}H_{260}N_{40}O_{38}S_4+H]^+$: 3706.8598, found 3706.8617. **LRMS:** (+ESI) *m/z* 1854.4 $[M+2H]^{2+}$, 1236.7 $[M+3H]^{3+}$, 927.7 $[M+4H]^{4+}$. **Analytical RP-HPLC:** $R_t$ = 20.3 min (0 to 70 vol.% MeCN in $H_2O$ with 0.1 vol.% TFA over 30 min, λ = 214 nm).

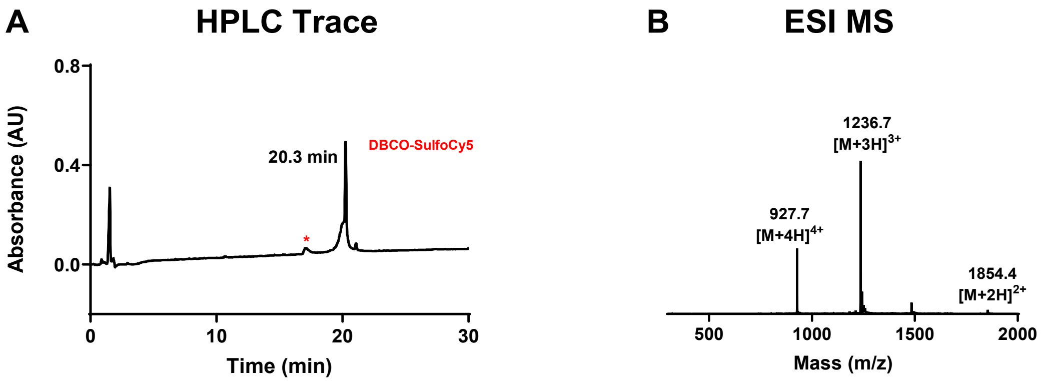

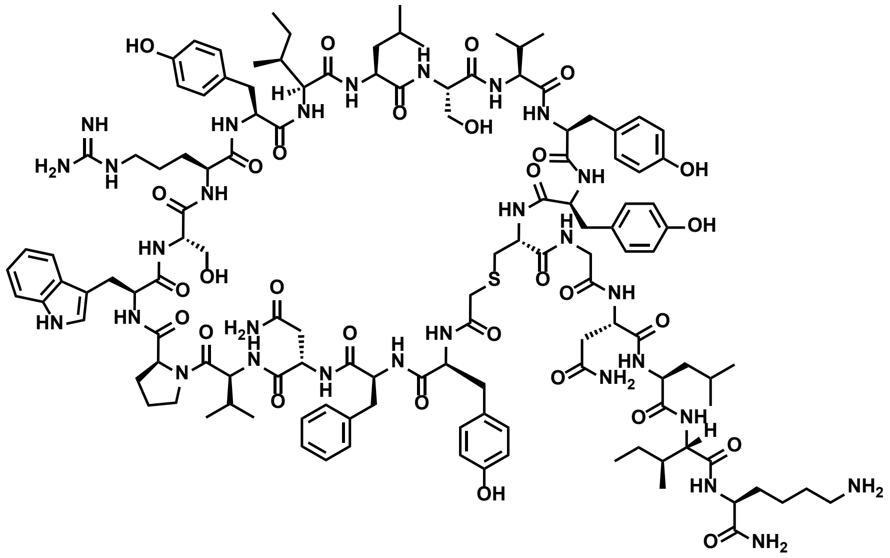

**A2*:** Rink amide resin (90 mg, 50 µmol, 0.56 mmol $g^{-1}$) was loaded with Fmoc-Lys(Boc)-OH according to general procedure **A**. The target peptide was then synthesised by iterative Fmoc-SPPS on the Syro I peptide synthesizer according to general procedure **A**, with chloroacetic acid as the final coupling step to generate the resin-bound sequence YFNVPWSRYILSVYYCGNLIK. The peptide was cleaved from resin by general procedure **C** and a portion (25 µmol) was cyclised according to general procedure **D**. Purification by preparative RP-HPLC (0 to 50 vol.% MeCN in $H_2O$ with 0.1 vol.% TFA over 50 min), followed by lyophilisation, afforded the pure peptide as a white fluffy solid (4.3 mg, 6%). **HRMS:** Calculated for $[C_{128}H_{181}N_{29}O_{30}S+H]^+$: 2637.3323, found 2637.3248. **LRMS:** (+ESI) *m/z* 1759.8 $[2M+3H]^{3+}$, 1320.0 $[M+2H]^{2+}$, 880.3 $[M+3H]^{3+}$. **Analytical RP-HPLC:** $R_t$ = 28.7 min (0 to 50 vol.% MeCN in $H_2O$ with 0.1 vol.% TFA over 30 min, λ = 214 nm).

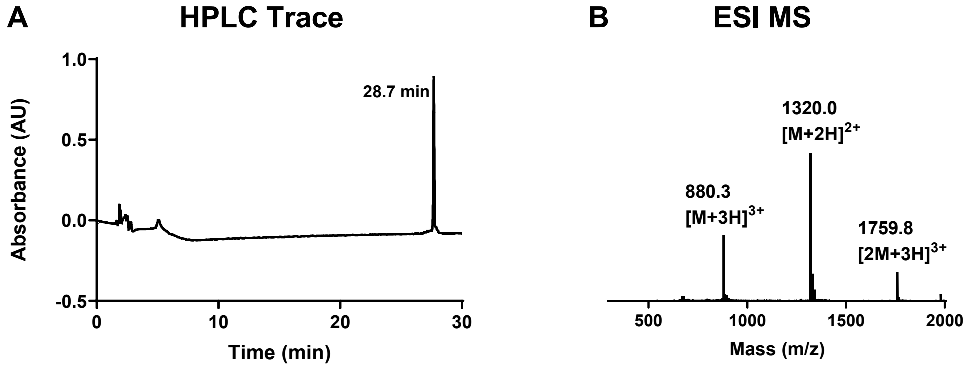

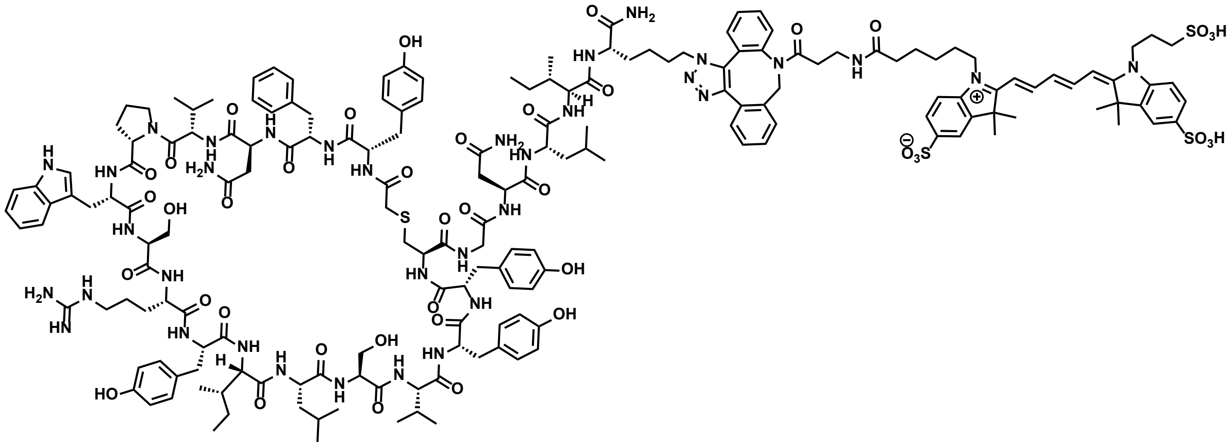

**Sulfo-Cy5-labelled A2*:** Rink amide resin (90 mg, 50 µmol, 0.56 mmol $g^{-1}$) was loaded with Fmoc-Lys*-OH according to general procedure **A** where Lys* refers to azidolysine. The target peptide was then synthesised by iterative Fmoc-SPPS on the Syro I peptide synthesizer according to general procedure **A**, with chloroacetic acid as the final coupling step to generate the resin-bound sequence YFNVPWSRYILSVYYCGNLIK*. The peptide was cleaved from resin by general procedure **C** and a portion (25 µmol) was cyclised according to general procedure **D**. Purification by preparative RP-HPLC (0 to 50 vol.% MeCN in $H_2O$ with 0.1 vol.% TFA over 50 min), followed by lyophilisation, afforded the pure peptide as a white fluffy solid (17.2 mg, 24%). DBCO-Sulfo-Cy5 was attached to the peptide via a SPAAC reaction (procedure **F**), purified by preparative RP-HPLC (0 to 50 vol.% MeCN in $H_2O$ with 0.1 vol.% TFA over 50 min) and lyophilised, isolating the final peptide as a blue fluffy solid (3.2 mg, 89%). **HRMS:** Calculated for $[C_{180}H_{235}N_{35}O_{41}S_4+H]^+$: not found. **LRMS:** (+ESI) *m/z* 1837.0 $[M+2H]^{2+}$, 1225.1 $[M+3H]^{3+}$. **Analytical RP-HPLC:** $R_t$ = 21.9 min (0 to 70 vol.% MeCN in $H_2O$ with 0.1 vol.% TFA over 30 min, λ = 214 nm).

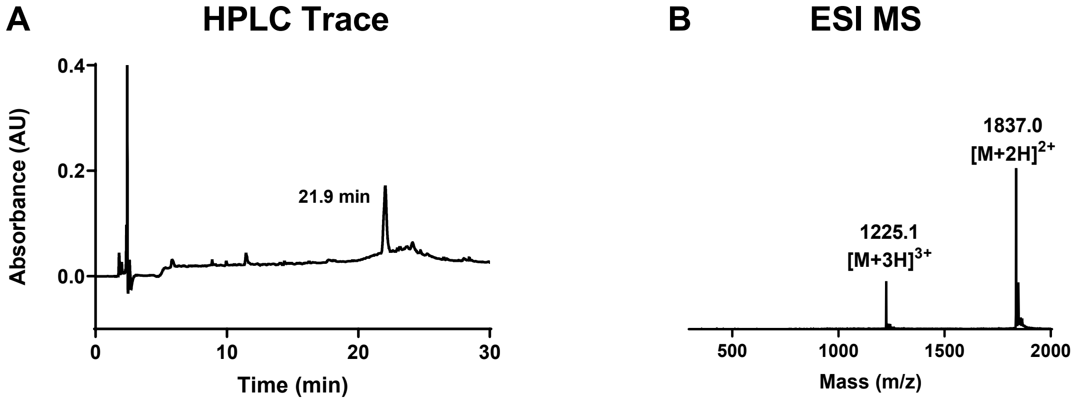

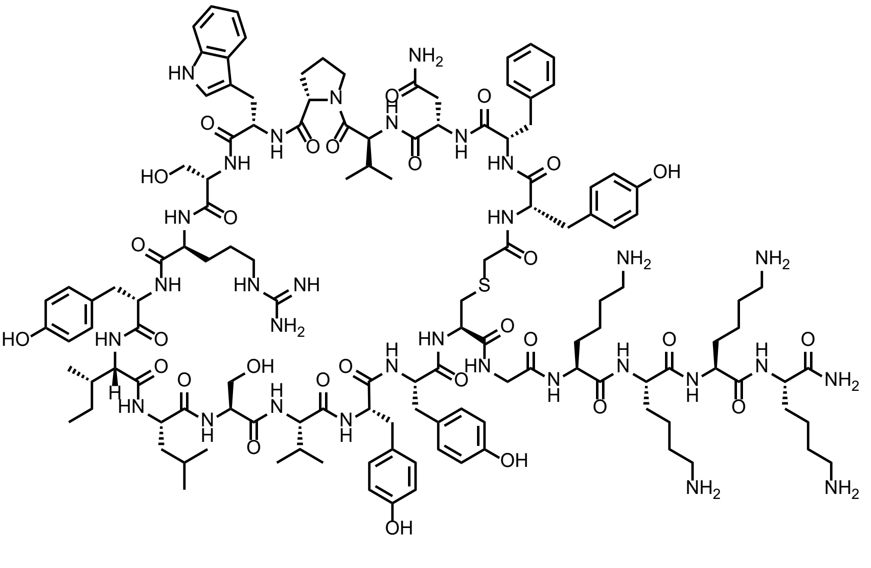

**A2Ktag:** Rink amide resin (90 mg, 50 µmol, 0.56 mmol $g^{-1}$) was loaded with Fmoc-Lys(Boc)-OH according to general procedure **A**. The target peptide was then synthesised by iterative Fmoc-SPPS on the Syro I peptide synthesizer according to general procedure **A**, with chloroacetic acid as the final coupling step to generate the resin-bound sequence YFNVPWSRYILSVYYCGKKKK. The peptide was cleaved from resin by general procedure **C** and a portion (5 µmol) was cyclised according to general procedure **D**. Purification by preparative RP-HPLC (0 to 60 vol.% MeCN in $H_2O$ with 0.1 vol.% TFA over 50 min), followed by lyophilisation, afforded the pure peptide as a white fluffy solid (3.2 mg, 20%). **HRMS:** Calculated for $[C_{130}H_{189}N_{31}O_{29}S+H]^+$: 2681.4061, found 2681.4090. **LRMS:** (+ESI) *m/z* 1787.6 $[2M+3H]^{3+}$, 1341.8 $[M+2H]^{2+}$. **Analytical RP-HPLC:** $R_t$ = 17.1 min (0 to 70 vol.% MeCN in $H_2O$ with 0.1 vol.% TFA over 30 min, λ = 214 nm).

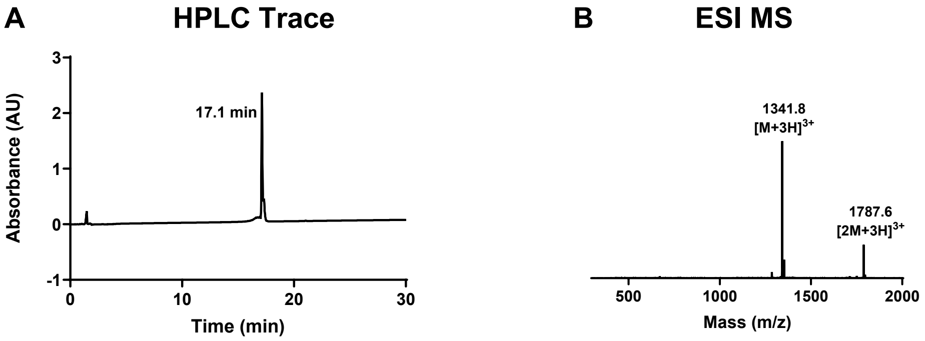

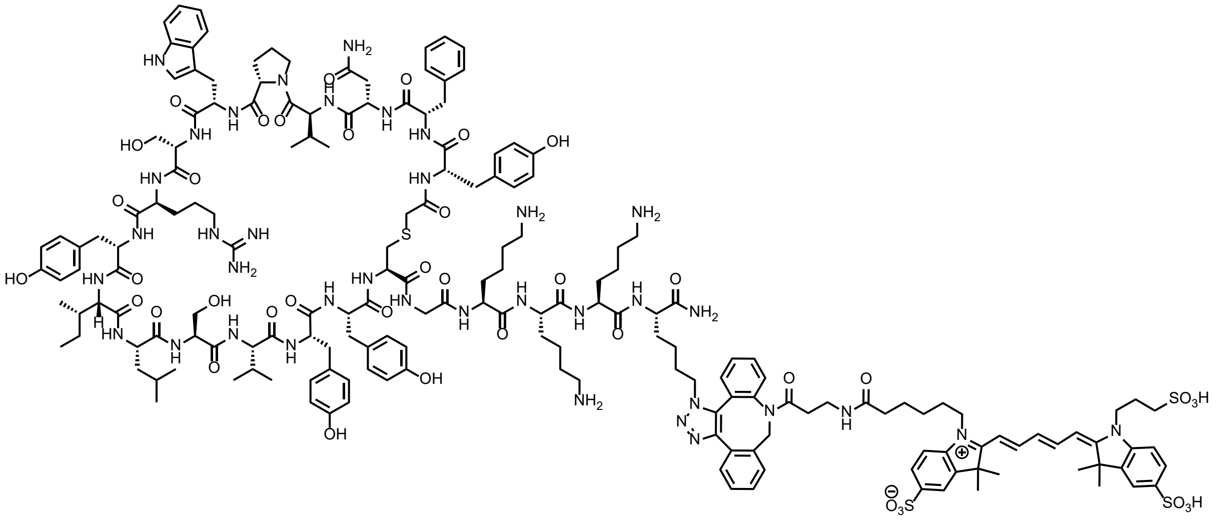

**Sulfo-Cy5-labelled A2Ktag:** Rink amide resin (90 mg, 50 µmol, 0.56 mmol g$^{-1}$) was loaded with Fmoc-Lys*-OH according to general procedure **A** where Lys* refers to azidolysine. The target peptide was then synthesised by iterative Fmoc-SPPS on the Syro **I** peptide synthesizer according to general procedure **A**, with chloroacetic acid as the final coupling step to generate the resin-bound sequence YFNVPWSRYILSVYYCGKKKK*. The peptide was cleaved from resin by general procedure **C** and a portion (5 µmol) was cyclised according to general procedure **D**. Purification by preparative RP-HPLC (0 to 60 vol.% MeCN in $H_2O$ with 0.1 vol.% TFA over 50 min), followed by lyophilisation, afforded the pure peptide as a white fluffy solid (5.1 mg, 32%). DBCO-Sulfo-Cy5 was attached to the peptide via a SPAAC reaction (procedure **F**) and purified through a silica plug. Following lyophilisation, the final peptide was isolated as a blue fluffy solid (1.8 mg, 73%). **HRMS:** Calculated for $[C_{182}H_{243}N_{37}O_{40}S_4+H]^+$: 3715.7074, found 3715.7055. **LRMS:** (+ESI) *m/z* 1239.7 $[M+3H]^{3+}$. **Analytical RP-HPLC:** $R_t$ = 19.1 min (0 to 70 vol.% MeCN in $H_2O$ with 0.1 vol.% TFA over 30 min, λ = 214 nm).

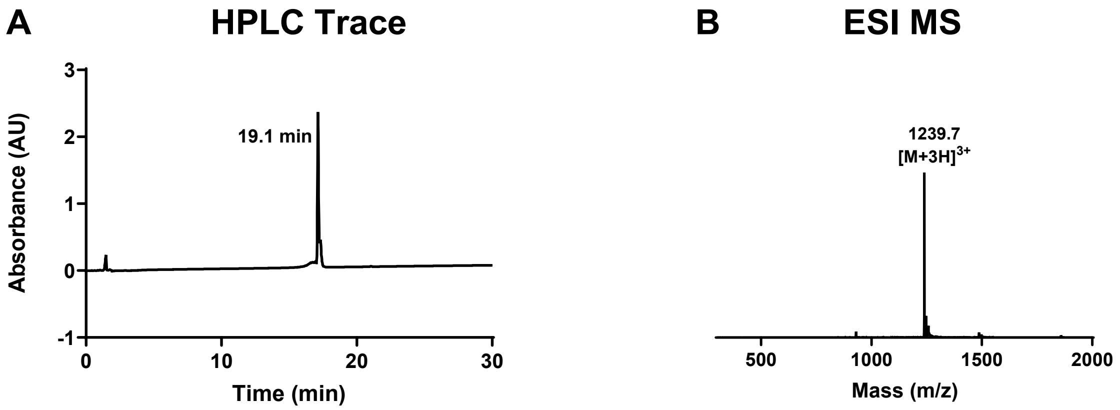

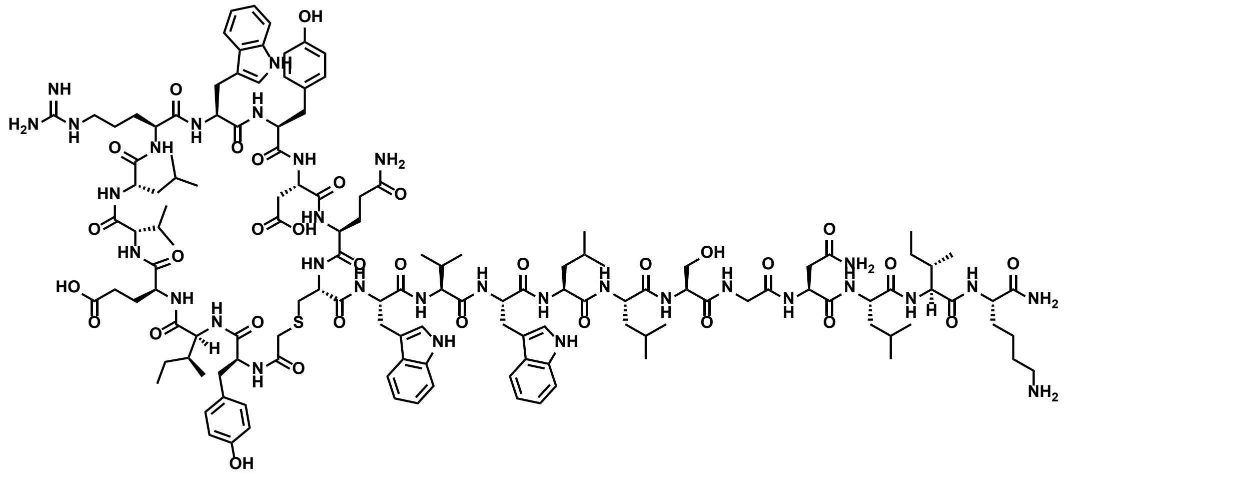

**A4*:** Rink amide resin (90 mg, 50 µmol, 0.56 mmol $g^{-1}$) was loaded with Fmoc-Lys(Boc)-OH according to general procedure **A**. The target peptide was then synthesised by iterative Fmoc-SPPS on the Syro I peptide synthesizer according to general procedure **A**, with chloroacetic acid as the final coupling step to generate the resin-bound sequence YIEVLRWYDQCWVWLLSGNLIK. The peptide was cleaved from resin by general procedure **C** and a portion (25 µmol) was cyclised according to general procedure **D**. Purification by preparative RP-HPLC (0 to 50 vol.% MeCN in $H_2O$ with 0.1 vol.% TFA over 50 min), followed by lyophilisation, afforded the pure peptide as a white fluffy solid (5.1 mg, 7%). **HRMS:** Calculated for $[C_{137}H_{198}N_{32}O_{32}S+Na]^+$: 2858.4463, found 2858.4550. **LRMS:** (+ESI) *m/z* 1892.8 $[2M+3H]^{3+}$, 1419.7 $[M+2H]^{2+}$. **Analytical RP-HPLC:** $R_t$ = 24.0 min (0 to 50 vol.% MeCN in $H_2O$ with 0.1 vol.% TFA over 30 min, λ = 214 nm).

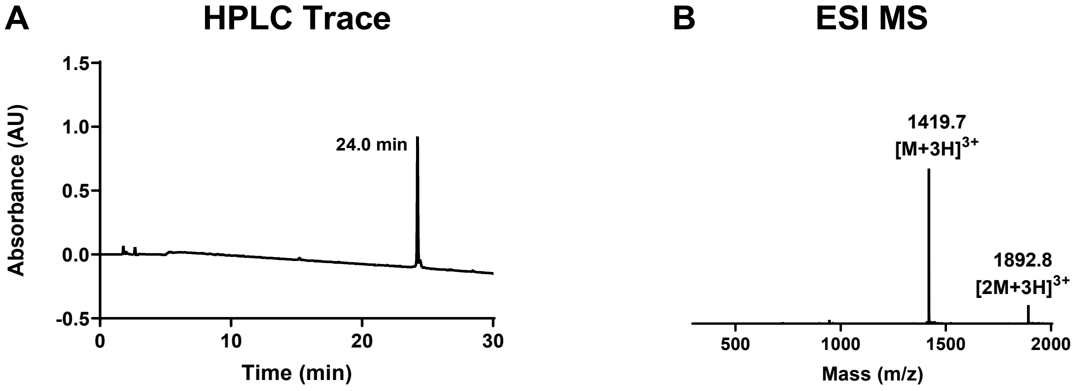

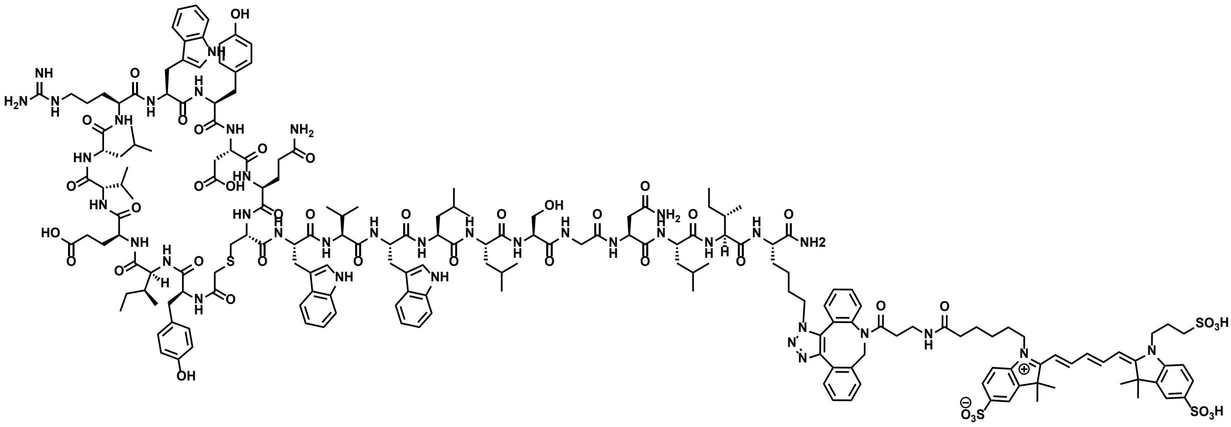

**Sulfo-Cy5-labelled A4*:** Rink amide resin (90 mg, 50 µmol, 0.56 mmol $g^{-1}$) was loaded with Fmoc-Lys*-OH according to general procedure **A** where Lys* refers to azidolysine. The target peptide was then synthesised by iterative Fmoc-SPPS on the Syro I peptide synthesizer according to general procedure **A**, with chloroacetic acid as the final coupling step to generate the resin-bound sequence YIEVLRWYDQCWVWLLSGNLIK*. The peptide was cleaved from resin by general procedure **C** and a portion (25 µmol) was cyclised according to general procedure **D**. Purification by preparative RP-HPLC (0 to 50 vol.% MeCN in $H_2O$ with 0.1 vol.% TFA over 50 min), followed by lyophilisation, afforded the pure peptide as a white fluffy solid (3.4 mg, 4%). DBCO-Sulfo-Cy5 was attached to the peptide via a SPAAC reaction (procedure **F**) and purified through a silica plug. Following lyophilisation, the final peptide was isolated as a blue fluffy solid (1.2 mg, 55%). **HRMS:** Calculated for $[C_{189}H_{252}N_{38}O_{43}S_4+H]^+$: not found. **LRMS:** (+ESI) *m/z* 1937.0 $[M+2H]^{2+}$, 1291.6 $[M+3H]^{3+}$, 969.0 $[M+4H]^{4+}$. **Analytical RP-HPLC:** $R_t$ = 24.6 min (0 to 70 vol.% MeCN in $H_2O$ with 0.1 vol.% TFA over 30 min, λ = 214 nm).

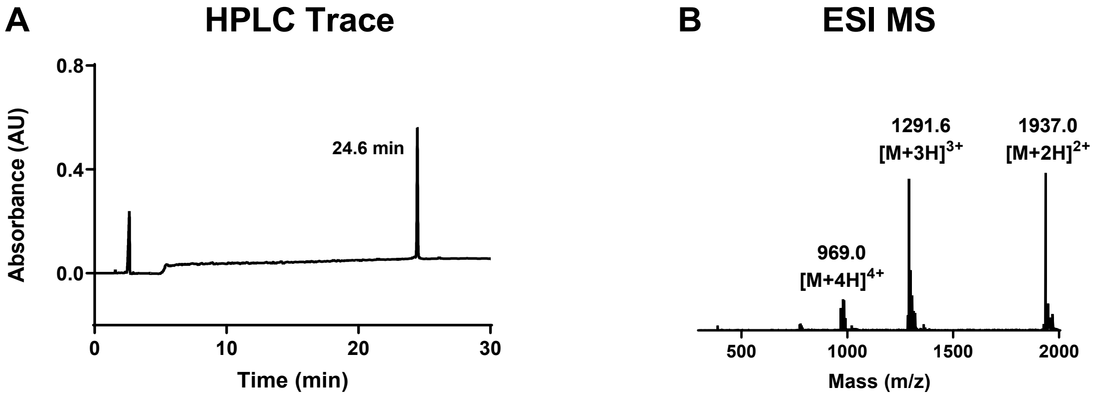

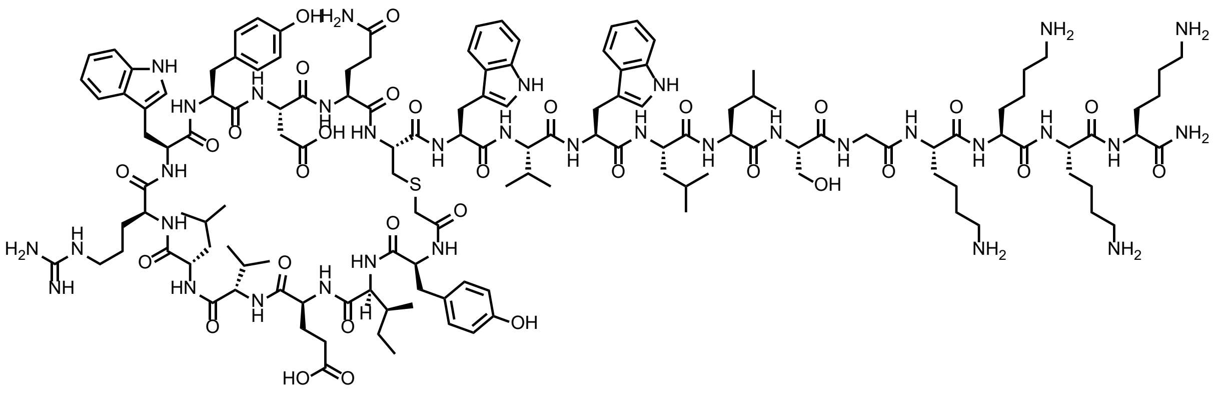

**A4Ktag:** Rink amide resin (90 mg, 50 µmol, 0.56 mmol $g^{-1}$) was loaded with Fmoc-Lys(Boc)-OH according to general procedure **A**. The target peptide was then synthesised by iterative Fmoc-SPPS on the Syro I peptide synthesizer according to general procedure **A**, with chloroacetic acid as the final coupling step to generate the resin-bound sequence YIEVLRWYDQCWVWLLSGKKKK. The peptide was cleaved from resin by general procedure **C** and a portion (5 µmol) was cyclised according to general procedure **D**. Purification by preparative RP-HPLC (0 to 60 vol.% MeCN in $H_2O$ with 0.1 vol.% TFA over 50 min), followed by lyophilisation, afforded the pure peptide as a white fluffy solid (1.0 mg, 6%). **HRMS:** Calculated for $[C_{139}H_{206}N_{34}O_{31}S+H]^+$: 2880.5382, found 2880.5386. **LRMS:** (+ESI) *m/z* 1921.8 $[2M+3H]^{3+}$, 1441.4 $[M+2H]^{2+}$. **Analytical RP-HPLC:** $R_t$ = 1688 min (0 to 70 vol.% MeCN in $H_2O$ with 0.1 vol.% TFA over 30 min, λ = 214 nm).

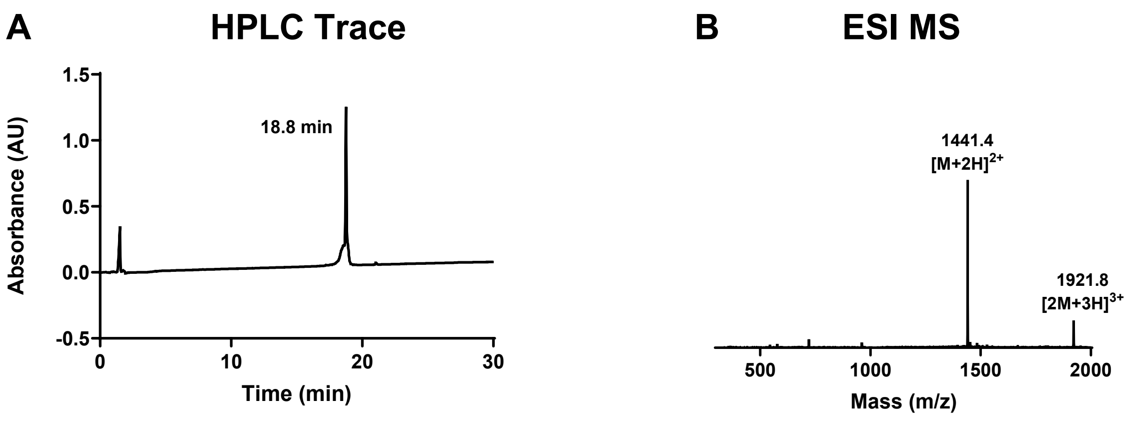

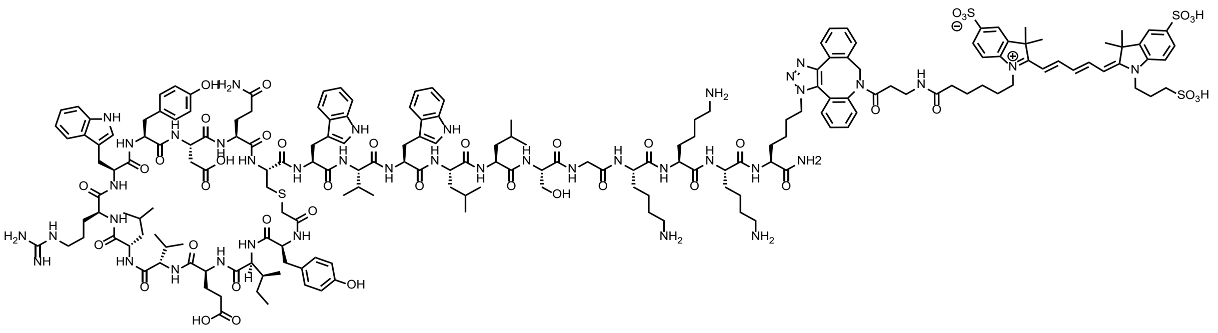

**Sulfo-Cy5-labelled A4Ktag:** Rink amide resin (90 mg, 50 µmol, 0.56 mmol g$^{-1}$) was loaded with Fmoc-Lys*-OH according to general procedure **A** where Lys* refers to azidolysine. The target peptide was then synthesised by iterative Fmoc-SPPS on the Syro I peptide synthesizer according to general procedure **A**, with chloroacetic acid as the final coupling step to generate the resin-bound sequence YIEVLRWYDQCWVWLLSGKKKK*. The peptide was cleaved from resin by general procedure **C** and a portion (5 µmol) was cyclised according to general procedure **D**. Purification by preparative RP-HPLC (0 to 60 vol.% MeCN in $H_2O$ with 0.1 vol.% TFA over 50 min), followed by lyophilisation, afforded the pure peptide as a white fluffy solid (0.9 mg, 6%). DBCO-Sulfo-Cy5 was attached to the peptide via a SPAAC reaction (procedure **F**) and purified through a silica plug. Following lyophilisation, the final peptide was isolated as a blue fluffy solid (0.7 mg, 60%). **HRMS:** Calculated for $[C_{191}H_{260}N_{40}O_{42}S_4+H]^+$: 3914.8394, found 3914.8394. **LRMS:** (+ESI) *m/z* 1306.1 $[M+3H]^{3+}$, 979.9 $[M+4H]^{4+}$. **Analytical RP-HPLC:** $R_t$ = 18.4 min (0 to 70 vol.% MeCN in $H_2O$ with 0.1 vol.% TFA over 30 min, λ = 214 nm).

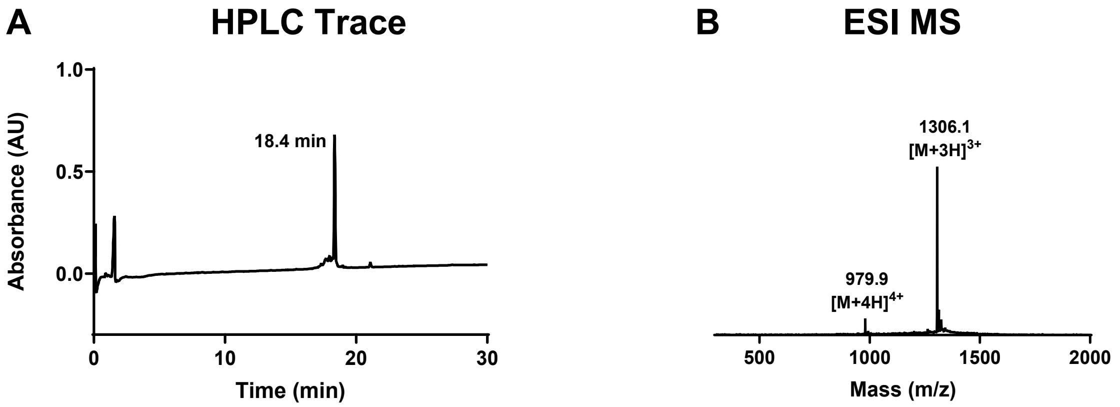
